# Genetic Perturbation Prediction Models Fail Out-of-Distribution: A Rigorous Benchmark and Systematic Evaluation

**DOI:** 10.64898/2026.10.16.759604

**Authors:** Kanupriya Tiwari, Gaurang Mahajan, Renuka Bellad, Nobal Kishor Dhruw, Abhishek Jha

## Abstract

Perturbation prediction models - which forecast transcriptional responses to gene inactivation - are increasingly proposed as *in silico* surrogates for genome-scale drug discovery screens. Yet, rigorous evaluation of whether these models genuinely generalize beyond their training distribution has been lacking. Existing benchmarks rely predominantly on random train-test splits, obscuring systematic failure modes that only emerge under realistic out-of-distribution (OOD) conditions. Here we introduce EL-PERTURB (Extrapolative Learning for PERTURBation Prediction), a benchmarking framework that stress-tests 20 perturbation prediction models spanning six architecture classes on both essential gene and genome-wide Perturb-seq screens, using a novel stratified OOD splitting strategy that partitions train and test sets into functionally distinct regions of transcriptional space. We find that model performance degrades substantially under OOD conditions, and that architectural complexity - including single-cell foundation model embeddings - confers no reliable advantage over well-tuned simpler baselines. EL-PERTURB establishes a rigorous benchmark for perturbation effect prediction across a gradient of generalization difficulty, providing a guide for building models that can genuinely serve as surrogates for wet-lab screening.

## 1 Introduction

Genome-scale genetic perturbation screens coupled with transcriptional readouts offer a mechanistically rich window into cellular processes, with transformative potential for target discovery and early drug development [1, 2]. Realizing this potential at scale requires computational models capable of predicting transcriptional responses to genetic perturbations - so-called *in silico* ‘virtual cell’ models - that can extrapolate reliably from a limited set of experimentally profiled knockdowns to the broader genome [3, 4]. The field has responded with a growing array of machine learning approaches spanning graph neural networks, deep learning frameworks, and foundation models fine-tuned for perturbation modeling [5, 6]. Yet the proliferation of proposed models has outpaced the development of rigorous benchmarks to assess whether these models genuinely deliver on their stated goals.

The core challenge is out-of-distribution (OOD) generalization. In a realistic target discovery setting, a model must predict responses to gene knockouts it has never seen - and often in cell lines it has never been trained on. This is a fundamentally different and harder problem than the interpolative task most models are actually evaluated on. The dominant evaluation practice in the field is random train-test splitting of perturbations, which inadvertently populates the test set with genes biologically similar to those in training, allowing models to succeed through nearest-neighbor interpolation rather than genuine extrapolation. Compounding this, most published benchmarks use generous training fractions of 50–80%, simulating data-rich conditions that bear little resemblance to the few-shot regime practitioners actually operate in [5, 7–9]. The result is a systematic overestimation of model capability - and a field that cannot distinguish genuine progress from increasingly sophisticated interpolation.

Several recent studies have begun to expose this gap. Ahlmann-Eltze et al. [10] compared seven models - five foundation models and two deep learning architectures - against linear baselines and found none outperformed simple alternatives, a finding independently corroborated by other studies showing that PCA-based regression and train-mean baselines match or exceed foundation model performance on perturbation tasks [9, 11]. While valuable, these studies share important limitations: they cover a narrow slice of the model landscape, do not employ stratified splitting strategies that enforce genuine distributional separation between train and test sets, and critically, offer no constructive prescription for what practitioners should do differently. The question of whether the problem lies in model architecture, training data design, or evaluation methodology itself has remained largely unanswered.

Here we introduce EL-PERTURB (Extrapolative Learning for PERTURBation prediction), a systematic benchmarking framework designed to answer that question (**Figure 1**). EL-PERTURB makes three distinct contributions. **First,** it provides a comprehensive head-to-head model comparison on single-gene perturbation prediction, evaluating 20 model configurations spanning six architecture classes - linear and kernel methods, multilayer neural networks, deep generative models, attention-based models, and graph neural networks - under a unified experimental protocol. **Second,** it applies a balanced eight-metric evaluation framework - combining transcriptome-level accuracy, perturbation-specific retrieval, and manifold preservation - designed to avoid the metric-specific biases that have distorted performance rankings in prior work. And **third,** it introduces a stratified OOD splitting strategy based on K-means clustering of ground truth transcriptional response profiles, which enforces functionally distinct train and test partitions and enables, for the first time, a reliable quantification of the OOD generalization gap across models. Together, our evaluation framework enables thorough analysis of genetic perturbation model performance across a gradient of generalization difficulty.

**Figure 1:**
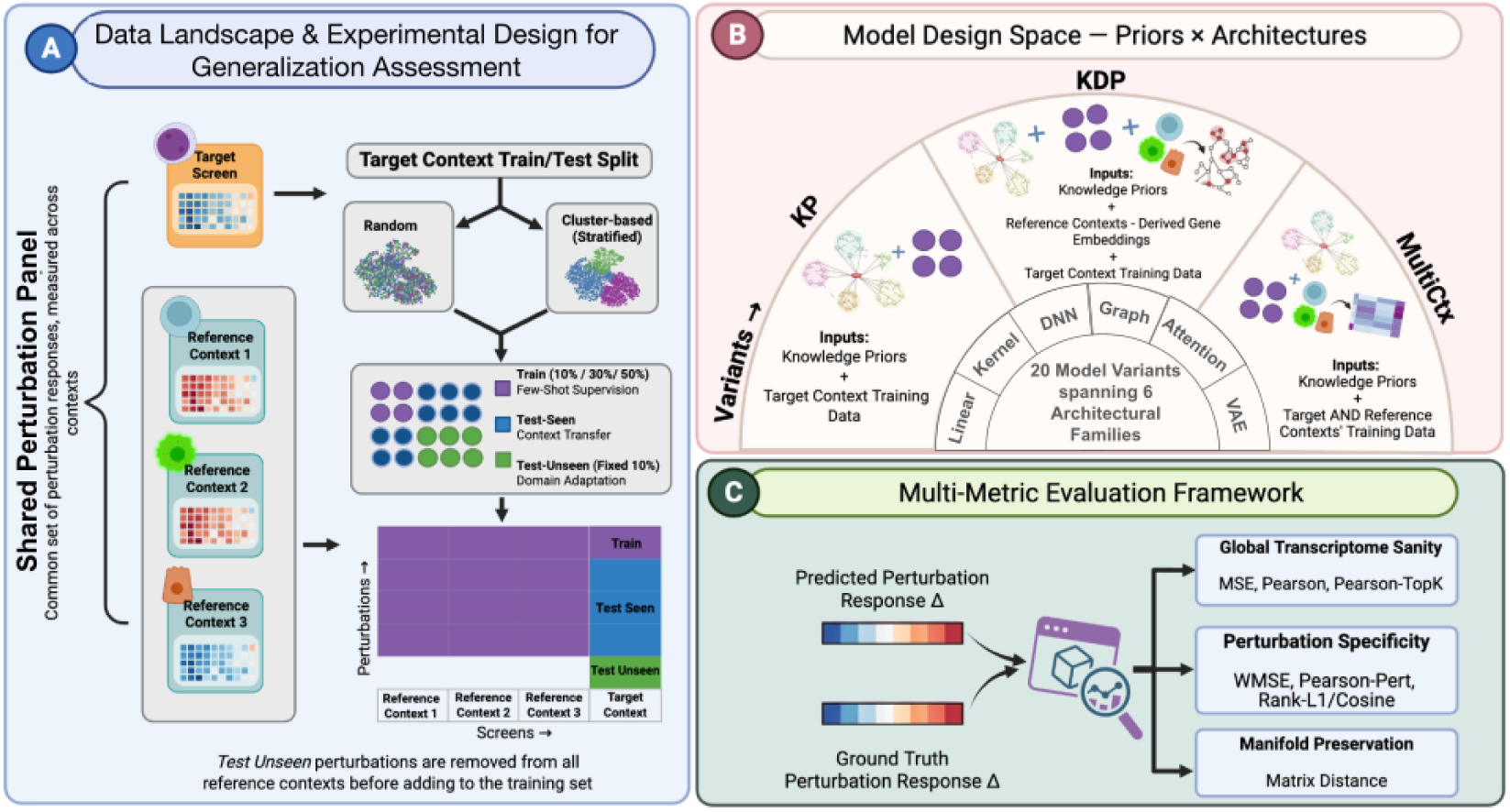
Integrated perturbation modeling framework and evaluation design. (A) To enable controlled evaluation of generalization under increasing distribution shift, a shared set of gene perturbations is profiled across multiple reference contexts and a target context. The target context is split into train (10%/30%/50% few-shot supervision), test-seen (context transfer), and test-unseen (domain adaptation, fixed at 10%) sets using either random or cluster-based stratified splits; test- unseen perturbations are removed from all reference contexts to prevent data leakage. (B) Twenty model variants are defined by crossing three data input strategies - knowledge priors with target context data (KP), knowledge priors with reference context-derived gene embeddings and target context data (KDP), and multi-context training integrating knowledge priors with target + reference context data (MultiCtx) - across six architectural classes (Linear, Kernel, DNN, Attention, GNN and VAE). (C) Predicted and ground-truth perturbation response deltas are compared using eight metrics organized into tiers of increasing biological stringency: global transcriptome-level agreement (MSE, Pearson, Pearson-TopK), perturbation specificity (WMSE, Pearson-Pert, Rank-L1/Cosine), and manifold preservation (Matrix Distance).

## 2 Related Work

### Evaluation metrics overestimate perturbation prediction performance

The metrics most commonly used to assess perturbation models have come under sustained scrutiny. Viñas Torné et al. [7] introduced Systema, demonstrating that standard metrics are susceptible to systematic variation - consistent transcriptional differences between perturbed and control cells arising from confounders rather than perturbation-specific biology. When this systematic component is removed, the apparent performance of existing methods drops substantially. Mejia et al. [8] traced the same problem to a specific mechanism: control-referenced delta metrics such as Pearson(Δ) are directly inflated by systematic bias between control and perturbed populations, and unweighted error metrics like MSE actively reward mode collapse - where models learn to predict the mean response across all perturbations regardless of the specific gene knocked out. This can be especially acute when modeling data from essential gene perturbation screens, which are expected to converge on common cellular stress response pathways. Their proposed remedies - DEG-weighted metrics, perturbed- mean referencing, and positive/negative baseline calibration - substantially reorder published method rankings. The PerturBench study [5] reported complementary findings, identifying mode collapse in published models and demonstrating the importance of rank-based metrics that test perturbation discriminability rather than aggregate regression accuracy. EL-PERTURB directly addresses this by adopting a balanced eight-metric framework that incorporates DEG-weighted, rank-based, and manifold-preservation metrics alongside standard correlations, explicitly designed to penalize mode collapse and avoid metric-specific ranking distortions.

### Existing benchmarks do not probe the full spectrum of generalization difficulty

Most published evaluations of perturbation prediction models, including those for GEARS [12] and PRESAGE [13], rely on random train-test splits with generous training fractions in the range of 50–80%, inadvertently populating test sets with perturbations biologically similar to training examples and allowing models to succeed through interpolation rather than genuine extrapolation. Recent benchmarking efforts have highlighted this problem from different angles: several studies [9–11] found that simple linear baselines match or outperform more complex models and foundation models on perturbation tasks. On the train/test partition side, PerturBench [5] and scPerturBench [14] introduced structured evaluation scenarios including unseen perturbations and cell types. However, none of these works directly compare random versus cluster-based splitting within the same cellular context to quantify how splitting strategy affects model rankings and reported performance. EL- PERTURB fills this gap with a stratified OOD splitting strategy based on K-means clustering of transcriptional response profiles, enabling systematic measurement of the OOD generalization gap across 20 models and six architecture classes.

### The model landscape for genetic perturbation prediction remains incompletely characterized

Genetic perturbation outcome prediction has been approached through two largely separate paradigms: in-context learning from prior knowledge features (e.g., GEARS, PRESAGE, GenePert [15]), and context transfer from cross-cell-line Perturb-seq data (e.g., CPA [16], scGen [17]). Models that integrate both - using prior knowledge embeddings alongside multi-context training data - have received comparatively little systematic evaluation, despite their theoretical advantage in the few-shot regime. While some recent models like TxPert [18] and scLAMBDA [19] are designed to integrate prior-knowledge perturbation space with cross-context perturbational data, the impact of multi-context training on domain adaptation and unseen gene generalization has not been systematically evaluated in the respective original studies. Moreover, the relative value of perturbation representations derived from curated biological knowledge graphs versus those from large-scale transcriptomic foundation models (e.g., scGPT, Geneformer [20, 21]) has not been rigorously compared under a controlled protocol. EL-PERTURB directly addresses both gaps by evaluating an integrated modeling framework across six architecture classes, systematically varying input featurization between curated knowledge priors and foundation model embeddings, and measuring the combined impact of knowledge priors and cross-context data across multiple generalization axes.

## 3 Methods

### 3.1 Data

We used essential-gene CRISPRi Perturb-seq screens from four human cell lines (K562, HepG2, RPE1, Jurkat) [22, 23] and, to test generalizability, single-cell data from four genome-wide CRISPRi screens (K562, HCT116, HEK293, resting CD4+ T cells) [22, 24, 25]. After knockdown-efficiency and coverage filtering, normalization, and HVG selection, each essential-gene screen comprised 1060 shared single-gene perturbations over 1463 genes, and each genome-wide screen 3065 single-gene perturbations over 3202 genes. The prediction target was the pseudobulk delta-expression vector: the shift from the mean log-normalized profile of non-targeting control cells. Full preprocessing, QC thresholds, and feature-selection details are provided in SI - **Detailed Methods**.

### 3.2 Modeling Framework

Genetic perturbation effect prediction was framed as a supervised regression problem: given a representation *I_P_* of a perturbed gene *P* , a model predicts the delta-expression response Δ = *f* (*I_P_* ) in a target cell line. Input representations combined biological prior knowledge with Perturb-seq response profiles induced by the same gene measured in other (reference) cell lines.

We compared three integration strategies for inputs that form the core axes of our experimental design (Figure 1): **KP** (knowledge priors only; single-context training), **KDP** (knowledge priors augmented with cross-context response profiles as input features), and **MultiCtx** (joint training across target and reference contexts with cell-line identity as an additional input). Where required, all models were supplied the same knowledge embeddings from PRESAGE [13] - 40 heterogeneous sources (including GO, Reactome, STRING, DepMap, ESM, BioGPT, GenePT), each PCA-reduced to *d* = 128 and concatenated.

Six architecture classes spanning a wide range of model complexity were benchmarked: attention-based (PRESAGE; KP, KDP, MultiCtx), multilayer neural networks (MLP with FiLM context injection; KP, KDP, MultiCtx), deep generative models (scLAMBDA [19]; KP, MultiCtx), linear and kernel regression methods (ridge, kernel ridge, latent ridge; KP, KDP), K-nearest neighbors on the combined kernel (KP, KDP), and LLM- and graph-based models (LangPert [26], an LLM- Ridge variant, and GEARS [12]; KP). The complete list of model configurations and details on hyperparameter tuning are provided in SI Table S1 and SI - **Detailed Methods**.

### 3.3 Few-Shot OOD Evaluation Framework

We used a leave-one-context-out design in which each cell line served as the target context in turn. To emulate data scarcity, in-context training fractions were set at 10%, 30%, and 50% of target-screen perturbations. Held-out perturbations were split into a *seen* subset (withheld from the target context but profiled in reference contexts, probing cross-context transfer) and an *unseen* subset (10% of perturbations, removed from all contexts, probing extrapolation to novel biology).

We contrasted two splitting strategies: a *random split*, testing interpolation, and a *stratified OOD split*, in which K-means clusters of ground-truth response profiles were assigned wholesale to train or test so the two sets occupied functionally distinct regions of transcriptional space. The OOD generalization gap was quantified as the percentage change in performance between the two conditions. For each training fraction and splitting strategy, we generated three independent splits using different random seeds for K-means clustering. Complete data splitting details, including the dependence of the distribution shift on the choice of K, are shared in SI - **Detailed Methods**.

### 3.4 Metrics

Predictions were evaluated with a rounded set of eight metrics at three levels of biological specificity: transcriptome-level accuracy (MSE, Pearson), biologically significant response accuracy (Pearson over the top-50 DEGs per perturbation), and perturbation-specific discriminability (WMSE [8], Pearson-Pert, Rank-L1, Rank-Cosine, and a Matrix Distance capturing global manifold structure). The perturbed gene was excluded throughout so that evaluation reflects trans-effects ignoring the trivial knockdown-efficacy signal.

Raw metrics were converted to Scaled Scores against a Cell-Mean baseline (the mean training-perturbation response), the mode-collapse reference: 1.0 represents perfect prediction, 0.0 matches the baseline, and negative values are worse than it. Two meta-scores were computed separately on seen and unseen subsets: the *Global Score* (GS; mean of all eight scaled metrics) and the *Specificity Score* (SS; mean of the five perturbation-specific discriminability metrics). Metric definitions and scaling formulas are provided in SI - **Detailed Methods**.

## 4 Results

We performed a systematic head-to-head comparison of pseudobulk genetic perturbation response prediction across 20 model configurations spanning six architecture classes, evaluating each under the KP, KDP, and MultiCtx integration strategies defined in Methods. Results are reported for the K562 target cell line as the primary test case for both the essential gene and genome-wide datasets, with reference data drawn from the remaining cell lines in each dataset (results from a second HepG2 cell line are provided as Supporting Data). Every model was separately tuned via 5-fold cross-validation on in-context training data, and test performance is reported for the best hyperparameters obtained. Results are organized around five questions: (1) How do models perform under in-distribution evaluation across essential gene and genome-wide screens? (2) How does performance change under stratified OOD splitting? (3) What role does dataset composition play in determining the benefit of cross-context data? (4) Do foundation model embeddings improve over curated knowledge priors? (5) Are model rankings stable across evaluation metrics?

### 4.1 Complex models provide no consistent advantage under in-distribution evaluation, and multi-context training helps only when cross-context signal exists

We first evaluated model performance under random train/test splits - the dominant evaluation practice in the field - on both essential gene and genome-wide screens.

We consider the essential gene panel first (Figure 2). As expected, performance improves with in-context training fraction across all models (TF 10% → 50%). Integrating cross-context data with knowledge priors consistently improves performance on the seen test subset across all model types, confirming effective transfer of shared response signatures from reference to target cell lines. For the multi-context MultiCtx models (PRESAGE-MultiCtx, scLAMBDA-MultiCtx and DNN-MultiCtx), this benefit extends to the unseen test subset - perturbations absent from all training contexts - particularly at low training fractions (TF=10%), where the MultiCtx variant of PRESAGE achieves more than 2x the Global Score of its single-context counterparts. This suggests that multi-context training enables genuine domain adaptation: alignment of perturbation representations to shared response structures that improves extrapolation even to genes never experimentally profiled in any context.

**Figure 2:**
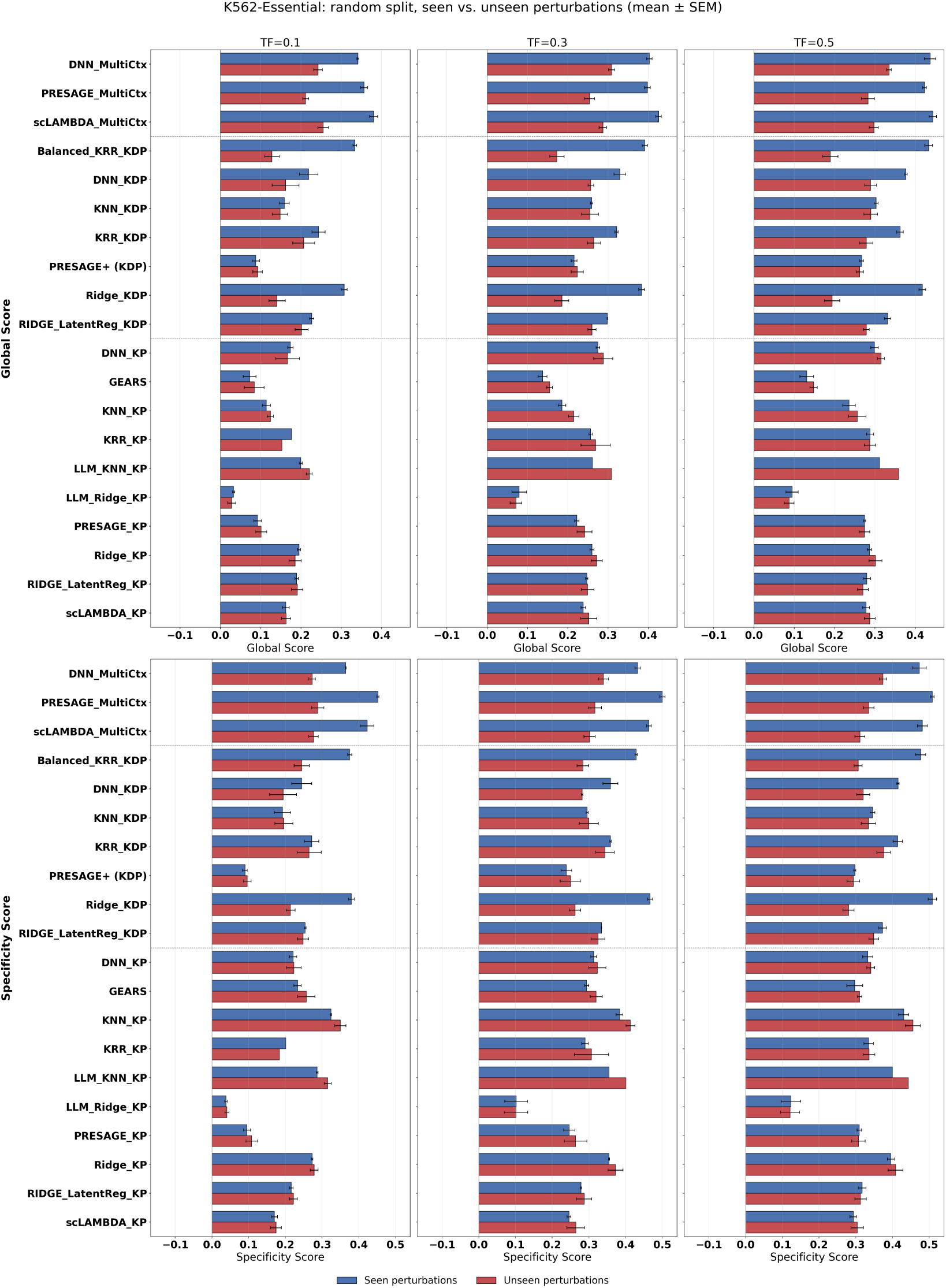
Model performance under random data splitting on seen vs. unseen perturbations (essential gene screen; target context = K562). Grouped bar plots comparing performance on seen (blue) and unseen (red) perturbations under the random split. Rows show the Global Score (top) and Specificity Score (bottom); columns show training fractions of 10%, 30% and 50%. Bars show the mean *±* SEM across random seeds.

However, the advantage of higher-capacity models is limited. Attention-based PRESAGE does not outperform MLP regression, likely reflecting the relatively small training set size ( few thousand perturbations). On the harder unseen test splits, simpler baselines - KRR-KDP, Latent-Ridge-KDP, and LangPert - perform on par with the deep learning models regardless of training fraction. No single model or architecture class dominates consistently across both seen and unseen test splits at all training fractions. Similar trends in model performance are apparent for the HepG2 essential genes panel (SI, Figure S1) as well.

The genome-wide screen results (Figure 3), on the other hand, tell a strikingly different story. Complex deep learning models are consistently and substantially outperformed by simpler baselines - not only on unseen perturbations but also on the seen subset, across all training fractions. Ridge regression, KNN regression, and LangPert are the top performers. LangPert - which relies exclusively on LLM-retrieved prior knowledge with no experimental data - outperforms multi-context deep learning models even on the seen test set (though a caveat being that frontier LLMs are pretrained on public biomedical literature and thus possibility of data leakage in this case cannot be ruled out; the alternative LLM-based method LLM-Ridge-KP which is less prone to such data leakage performs comparatively poorly). Integration of cross-context data with knowledge priors yields no consistent benefit in this setting, and overall Global/Specificity Scores are substantially lower than on the essential gene panel across the board.

**Figure 3:**
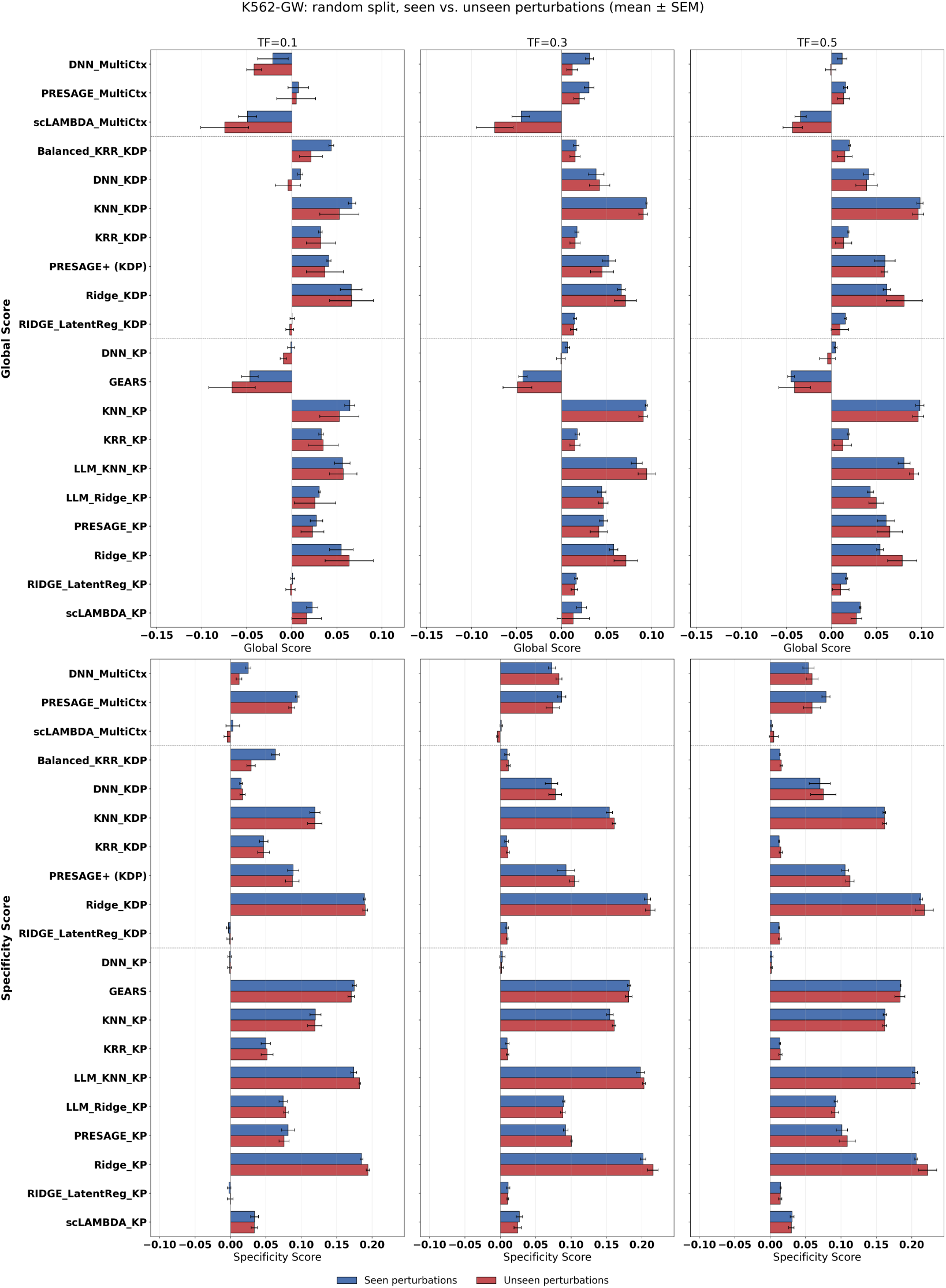
Model performance under random data splitting on seen vs. unseen perturbations (genome wide screen; target context = K562). Grouped bar plots comparing performance on seen (blue) and unseen (red) perturbations under the random split. Rows show the Global Score (top) and Specificity Score (bottom); columns show training fractions of 10%, 30% and 50%. Bars show the mean *±* SEM across random seeds.

### 4.2 All models fail under stratified OOD splitting - complexity does not help

Having established in-distribution baselines, we next applied our stratified OOD splitting strategy, which partitions perturbations into functionally distinct train and test clusters based on K-means clustering of ground truth response profiles. This is the key stress test of EL-PERTURB: it forces models to extrapolate to perturbations in transcriptionally novel regions of response space rather than interpolate within familiar neighborhoods.

In the case of the K562 essential genes screen, performance degrades substantially under stratified splits relative to random splits, particularly on the test-unseen partition (Figure 4). The OOD generalization gap - defined as the percentage change in score between random and stratified conditions on the unseen test set - is largely negative, with gaps ranging from moderate (approximately *−*10%) to severe (exceeding *−*40% for several models). The gap is in fact more acute when performance is assessed in terms of the Specificity Score. Increasing in-context training data marginally reduces the gap but does not close it. Critically, there is no consistent relationship between model complexity and OOD robustness: simpler baselines are no more vulnerable than deep learning models, and complex architectures provide no particular advantage. The impact of distribution shift is similarly visible in the results for the HepG2 essential genes panel (SI, Figure S2).

**Figure 4:**
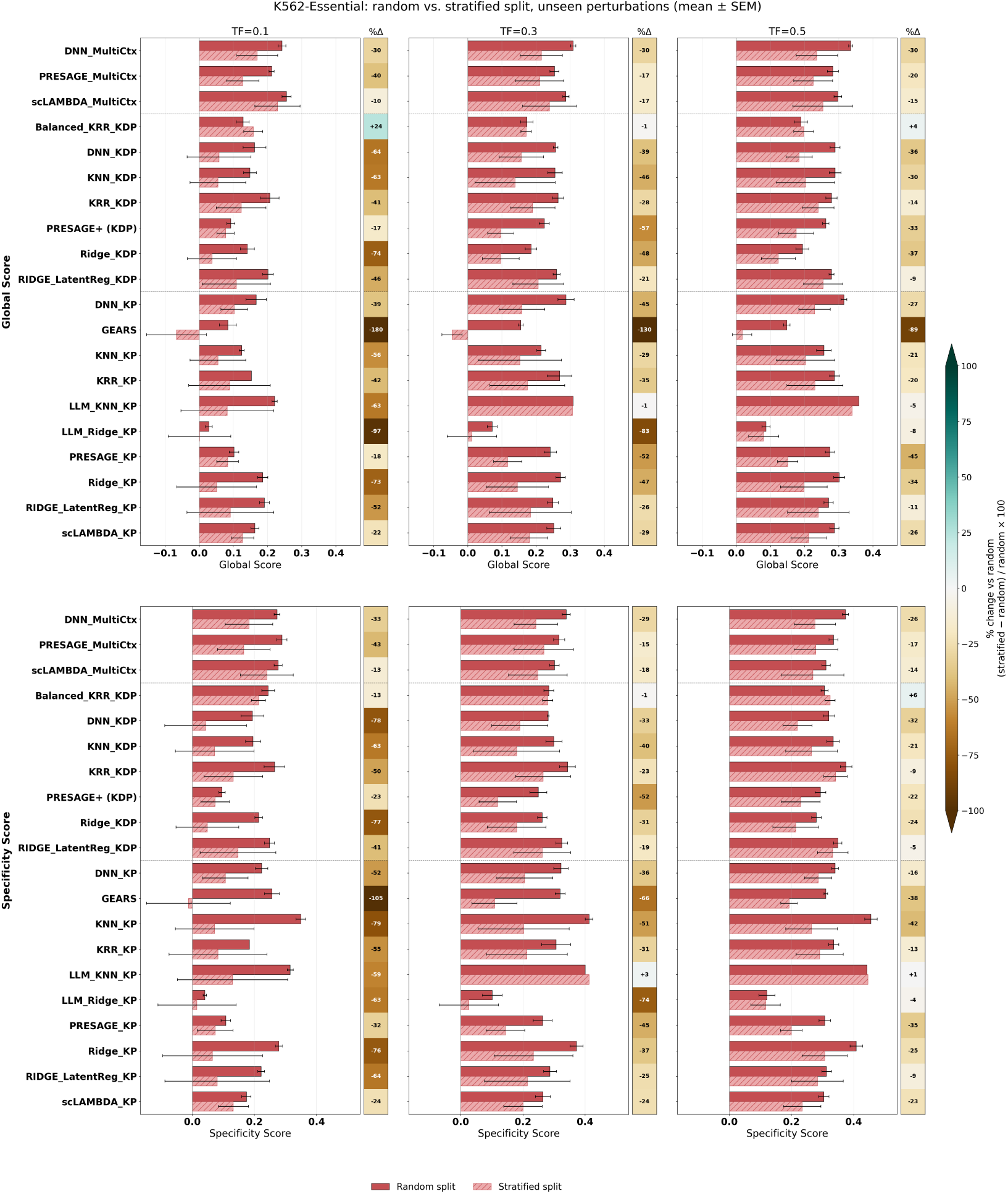
OOD generalization gap across models for essential gene screens (target context = K562). Grouped bar plots comparing performance on unseen perturbations under random (solid red) and stratified (light red, hatched) splits. Rows show the Global Score (top) and Specificity Score (bottom); columns show training fractions of 10%, 30% and 50%. Bars show the mean *±* SEM across random seeds. The column of squares to the right of each panel (%Δ) gives the relative change under the stratified split, (stratified *−* random) / random *×* 100, for each model; brown indicates lower and teal higher performance than under the random split, with the colour scale capped at *±*100%. Any cells marked n/a are omitted because the random-split score is below 0.01, where a relative change is not meaningful.

On the K562 genome-wide screen (Figure 5), all models effectively fail under both random and stratified splits, with Global as well as Specificity Scores consistently in a low range on the challenging test-unseen partition indicating little predictive signal beyond the Cell-Mean baseline. The test performance on stratified splits is by and large lower than on the random splits, with the OOD gap being relatively more pronounced for Specificity Scores. There are a few stray instances which break the general pattern, but this is not consistent across the evaluated training sizes, and is offset by the poor absolute performance of these models on the genome-wide screen, besides partly accounted by the higher variability in the test performance across seeds under stratified splitting making the random versus stratified differences noisier in general. Further, the relative ranking of models bears little resemblance to results on the essential gene panel - underscoring that performance on one dataset cannot be extrapolated to another (SI, Figures S3 and S4). No deep learning variant is consistently performant across the board - GEARS fares poorly on the multi-metric evaluation, PRESAGE and scLAMBDA (MultiCtx variants) do relatively better on essential gene panels but this does not carry over to the GW screen.

**Figure 5:**
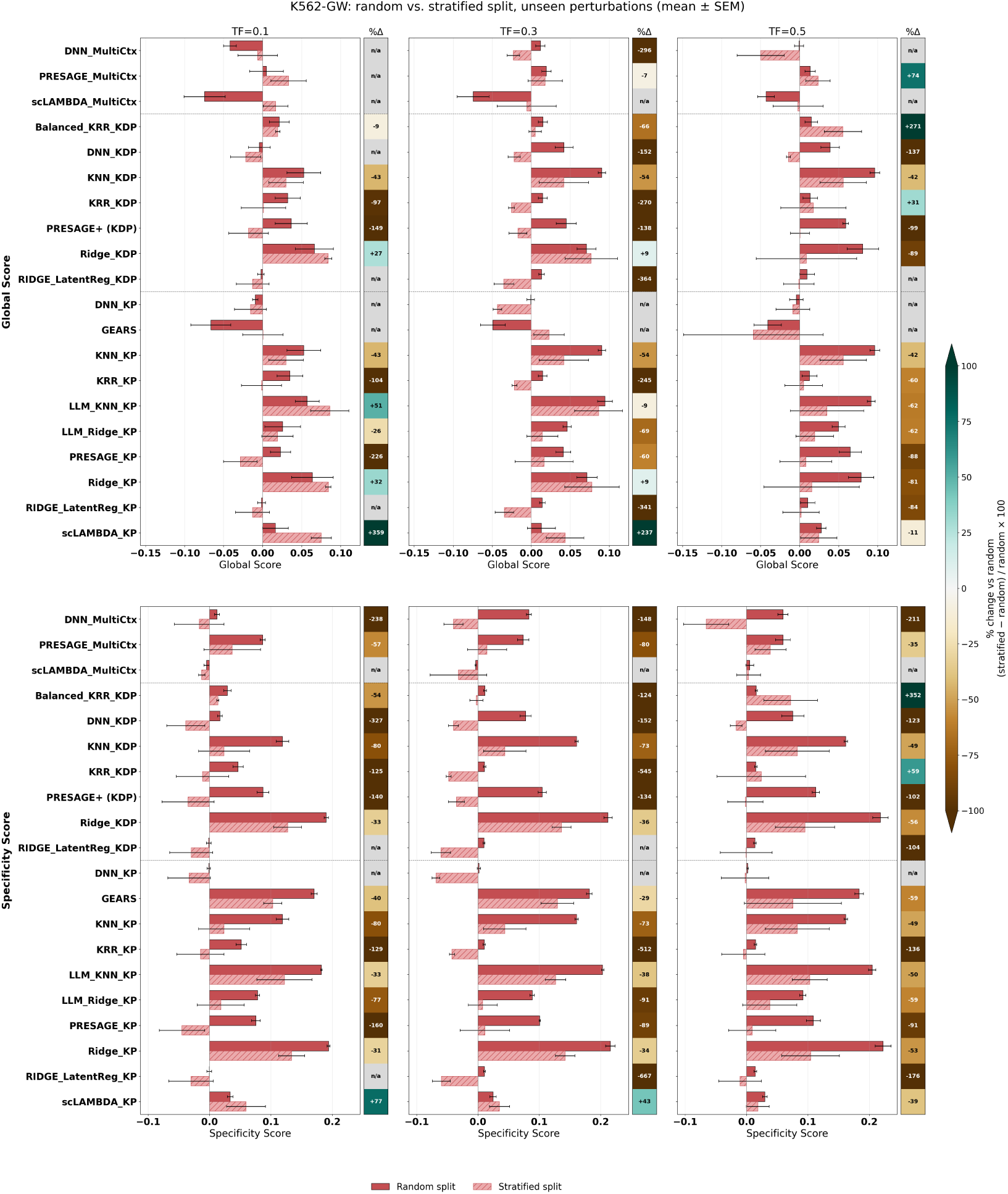
OOD generalization gap across models for genome-wide screen (target context = K562). Same format as Figure 4.

### 4.3 Cross-context transferability depends on shared response structure

The contrast between essential gene and genome-wide results raises a natural question: why does multi-context training help in one setting but not the other? To answer this, we characterized the diversity of each dataset through a set of heuristics capturing both cross-context transferable response structure and in-context response heterogeneity (Figure 6 and SI Table S2).

**Figure 6:**
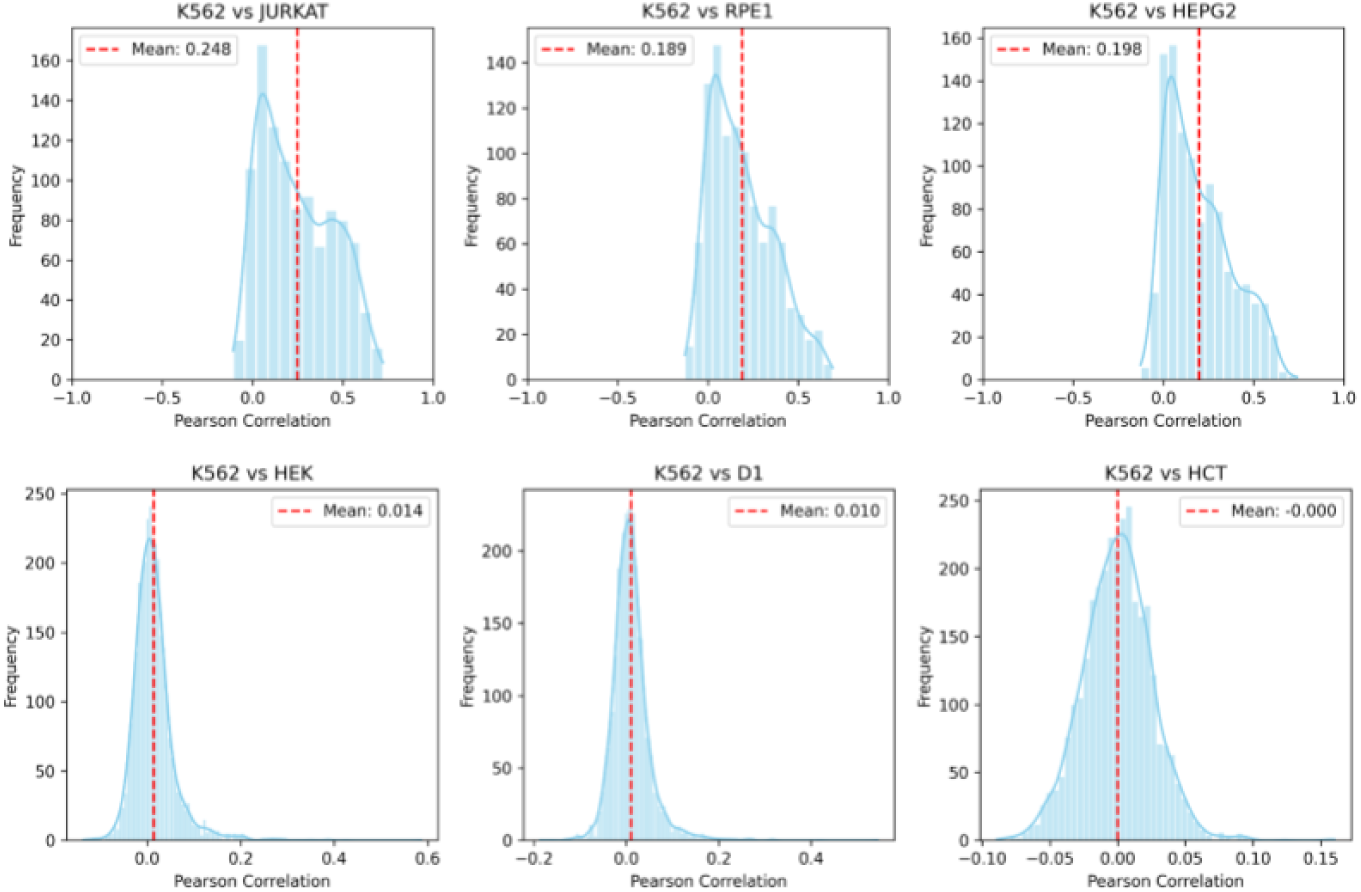
Cross-context correlation in pseudobulk logFC perturbation response profiles between target (K562) and reference contexts in essential genes-only (top row) and genome-wide (bottom row) experiments.

The data indicate that CRISPR knockdowns of essential genes produce relatively more correlated transcriptional response signatures across the cell lines considered, providing an informative shared signal for cross-context transfer. The genome-wide perturbation screens, by contrast, span a far more heterogeneous phenotypic space, with substantially weaker cross-context as well as in-context correlations. This explains why multi-context training benefits prediction in the essential gene setting but fails in the genome-wide case: interpolative models reliant on memorization can only leverage cross-context data when that data is correlated with the target context response.

### 4.4 Foundation model embeddings do not improve over curated knowledge priors

Next, we evaluated gene embeddings from two single-cell transcriptomics foundation models: pre-trained scGPT [20] and Geneformer [21]. These include direct gene representations, as well as *in silico* perturbation (ISP) embeddings generated using the approach described in the Geneformer paper which carry a causal flavor by modeling perturbation-induced cell state shifts in the foundation model’s representation space.

In head-to-head comparison across both MLP and Ridge Regression models on the K562 essential gene screen (Figure 7), foundation model embeddings substantially underperform curated knowledge-derived embeddings (the 40-source PRESAGE embedding set). Further, when FM-derived perturbation embeddings are combined with the curated embeddings, they provide no additive improvement - and in some configurations slightly degrade performance. This pattern holds across both single-context and multi-context settings and across all training fractions. These results suggest that the single-cell FM embedding space, as currently formulated, does not encode information relevant to perturbation response prediction that is not already captured by curated knowledge sources.

**Figure 7:**
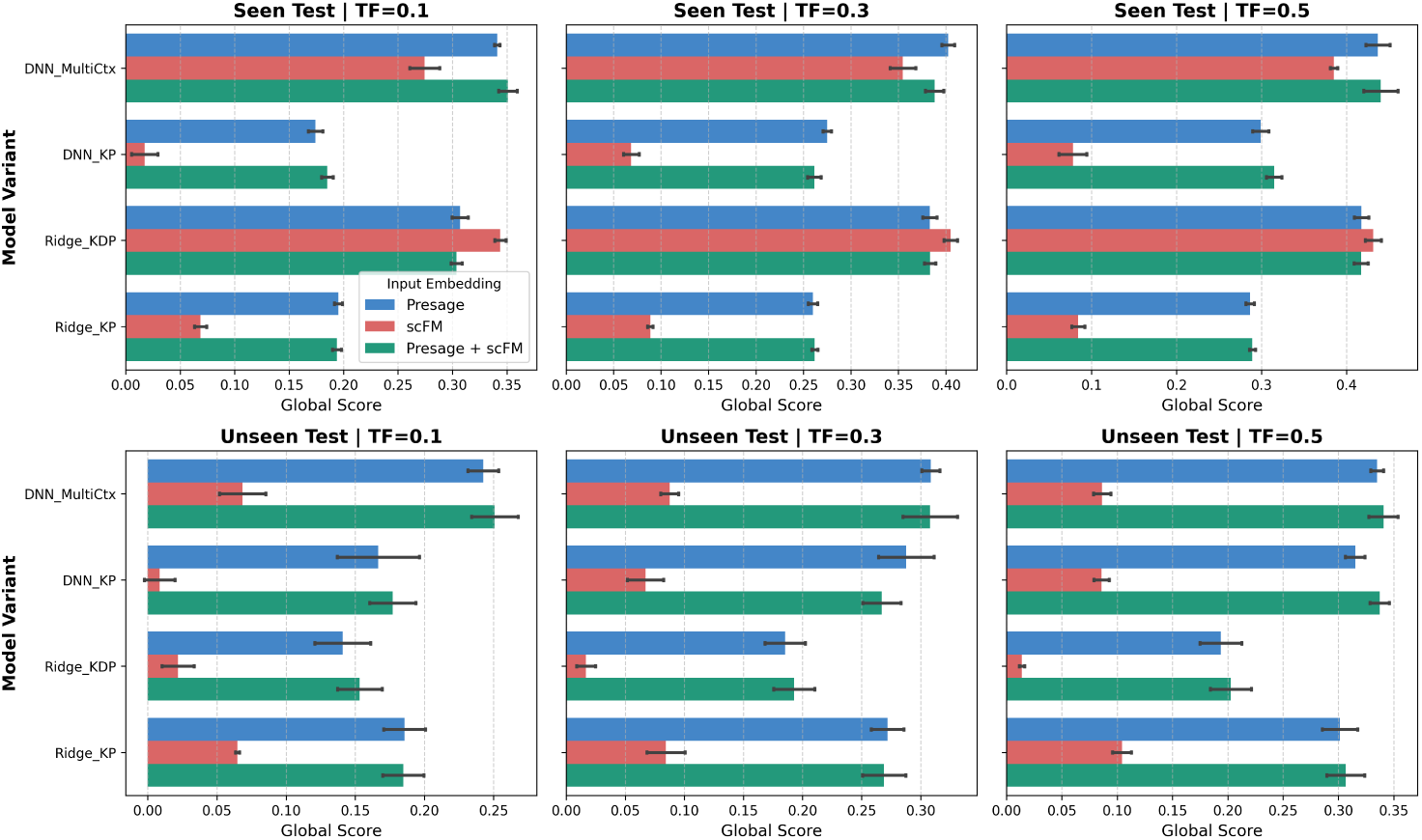
Single-cell foundation model embeddings do not improve perturbation response prediction over curated knowledge-derived features. Barplots comparing Global Score across gene embedding strategies (FM only, curated only, FM + curated) for DNN and Ridge Regression on the K562 essential gene panel.

### 4.5 Model rankings are metric-dependent, supporting the need for multi-metric evaluation

*Individual metric contributions to the Global and Specificity Scores differ between essential-gene and genome-wide screens:*

Figures 8 and 9 report, for each evaluation condition (random/stratified *×* seen/unseen), the spread of each scaled metric, its rank correlation with each aggregated score, and its share of each score’s variance across models.

**Figure 8:**
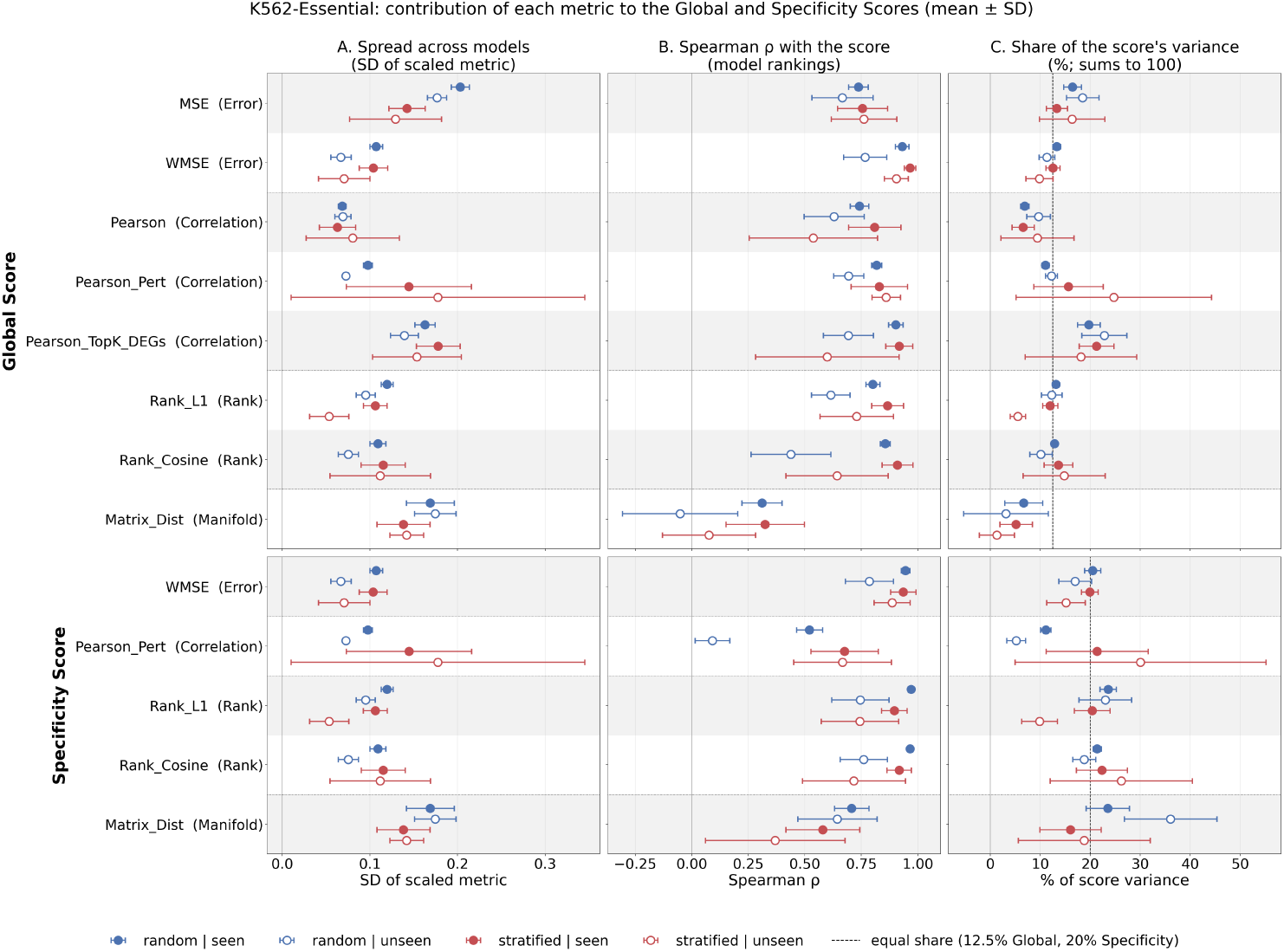
Effective contribution of each metric to the Global and Specificity Scores (K562- Essential). Rows show the Global Score (top; 8 metrics) and Specificity Score (bottom; 5 metrics). For each evaluation condition (blue = random, red = stratified split; filled = seen, open = unseen perturbations), we show (A) the spread of each scaled metric across models (standard deviation), (B) the Spearman rank correlation between each metric and the score across models (+1 = the metric ranks models in the same order as the score), and (C) each metric’s share of the score’s variance across models. Shares sum to 100% and can be negative when a metric works against the score; the dashed line marks an equal share (12.5% for the Global Score, 20% for the Specificity Score). Points show the mean *±* SD over the nine runs per condition (three training fractions three random seeds). Dotted lines separate metric families.

**Figure 9:**
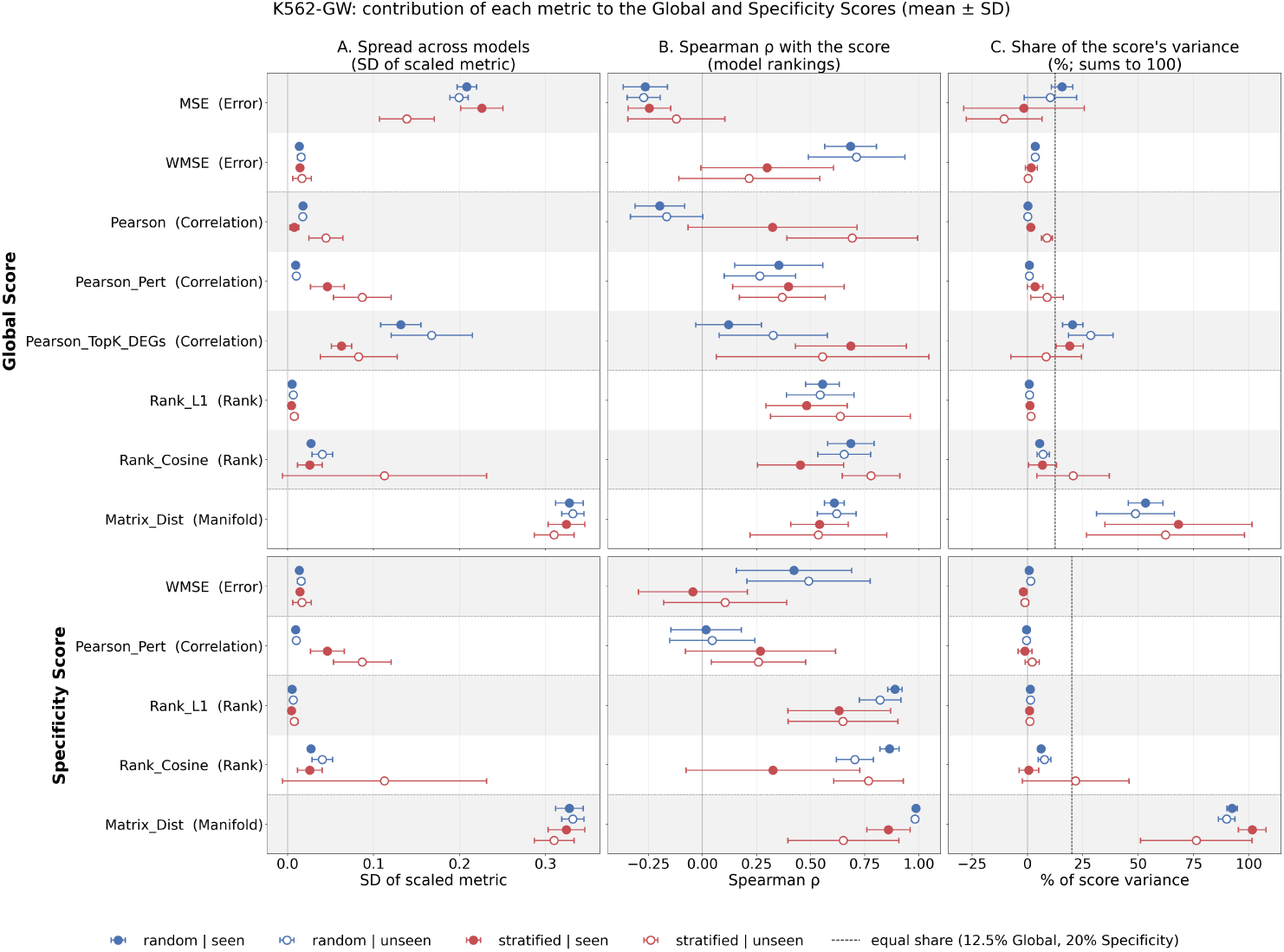
Effective contribution of each metric to the Global and Specificity Scores (K562-GW).

On K562-Essential (Figure 8), most metrics contribute close to their equal share. Outside stratified/unseen, MSE, WMSE, Pearson-Pert and the two rank scores each account for 10–18% of GS variance (equal share: 12.5%). Pearson-TopK-DEGs contributes more (20–23%), whereas Pearson (7–10%) and Matrix Distance (3–7%) contribute less: Matrix Distance varies across models as much as Pearson-TopK-DEGs but ranks them differently from the other metrics. For SS (equal share: 20%), WMSE and the two rank scores contribute 17–24% and Matrix Distance 16–36%, while Pearson-Pert rises from 5–11% under random splits to 21% under stratified/seen. Stratified/unseen extends this shift towards Pearson-Pert (25 *±* 20% of GS, 30 *±* 25% of SS) at the expense of Rank-L1 (5 *±* 2% and 10 *±* 4%), but only the Rank-L1 decrease is consistent across seeds. Model rankings in this condition are far less reproducible: on the Essential datasets, between-seed Spearman *ρ* is 0.26–0.65, compared with 0.77–0.98 in the other conditions (SI Table S3). HepG2-Essential shows a similar pattern (SI, Figure S5).

K562-GW behaves differently (Figure 9). Matrix Distance accounts for 49–68% of GS and 76–100% of SS variance in every condition; under random splits, Pearson-TopK-DEGs (20–29%) and MSE (10–16%) are the only other substantial contributors to GS. This reflects how little models improve on the baseline (best GS *≤* 0.09, compared with 0.24–0.42 on K562-Essential): most scaled metrics barely separate models, whereas Matrix Distance varies more than ten times as much as Pearson, WMSE or Rank-L1. Although reproducible between seeds under random splits (*ρ* = 0.84–0.98), the GW rankings largely reflect this single metric.

*Different metric families favor different architectures:*

Since K562-GW models barely improve on the baseline and their aggregate rankings seem to be largely driven by Matrix Distance alone, we examine how architecture interacts with metric choice mainly on K562-Essential results, where all metric families contribute meaningfully. Averaging ranks within four metric groups (error, correlation, rank and manifold) shows consistent architecture profiles on K562-ESS in every evaluation condition (Figure 10). DNN and VAE models (scLAMBDA) rank among the best on error and correlation metrics but in the lower half on Matrix Distance. KNN-based models show the opposite profile, weak on error and correlation but strong on rank and manifold metrics. GEARS is the extreme case, ranking last or near-last on error and correlation yet first on Matrix Distance, while linear models stay near the middle across all metric groups.

**Figure 10:**
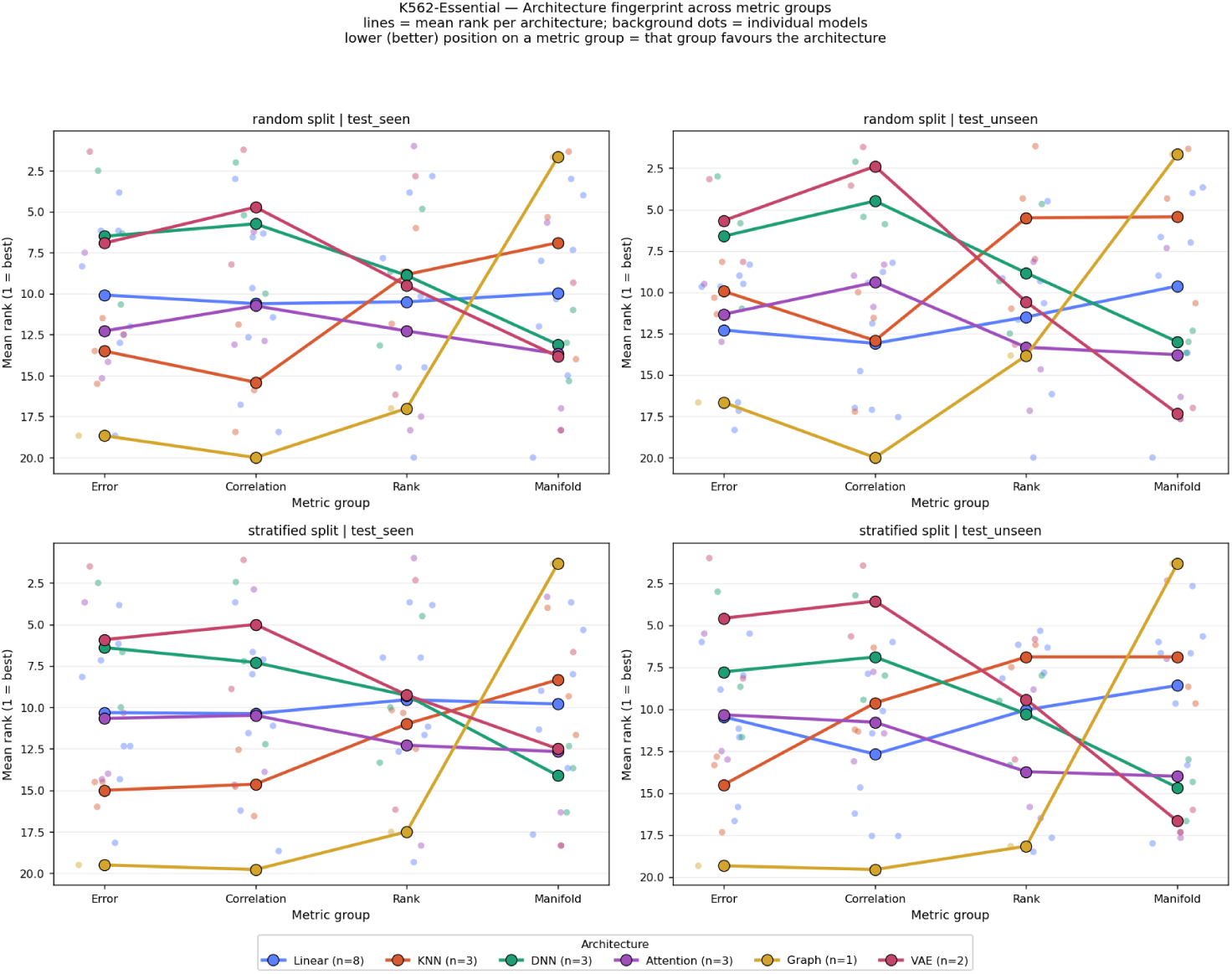
Architecture-specific performance profiles across metric groups (K562-Essential). For each evaluation condition (panels), models were ranked on each metric (averaged over random seeds; rank 1 = best), and ranks were averaged across training fractions and then within four metric groups: Error (MSE, WMSE), Correlation (Pearson, Pearson-Pert, Pearson-TopK-DEGs), Rank (Rank-L1, Rank-Cosine) and Manifold (Matrix Distance). Lines show the mean rank per architecture class, and background points show individual models; n gives the number of models per class. Higher positions indicate better ranks.

Figure 11 further depicts the ranking differences between the models based on the GS and SS scores. For most models, the two ranks agree within *±*3 positions. The systematic exceptions follow the architecture profiles. Models strong on rank and manifold metrics rank higher under SS, where Matrix Distance carries more weight: GEARS gains up to 11 positions, and KNN_KP gains 10–15 positions under random splits. Models strong on error and correlation metrics rank higher under GS: scLAMBDA gains up to 8 positions, and DNN models up to 6. Notably, PRESAGE-MultiCtx is among the most consistent models (|gap| *≤* 2 in all configurations, Spearman *ρ ≈* 0.95), consistent with its balanced profile observed in Figure 10.

**Figure 11:**
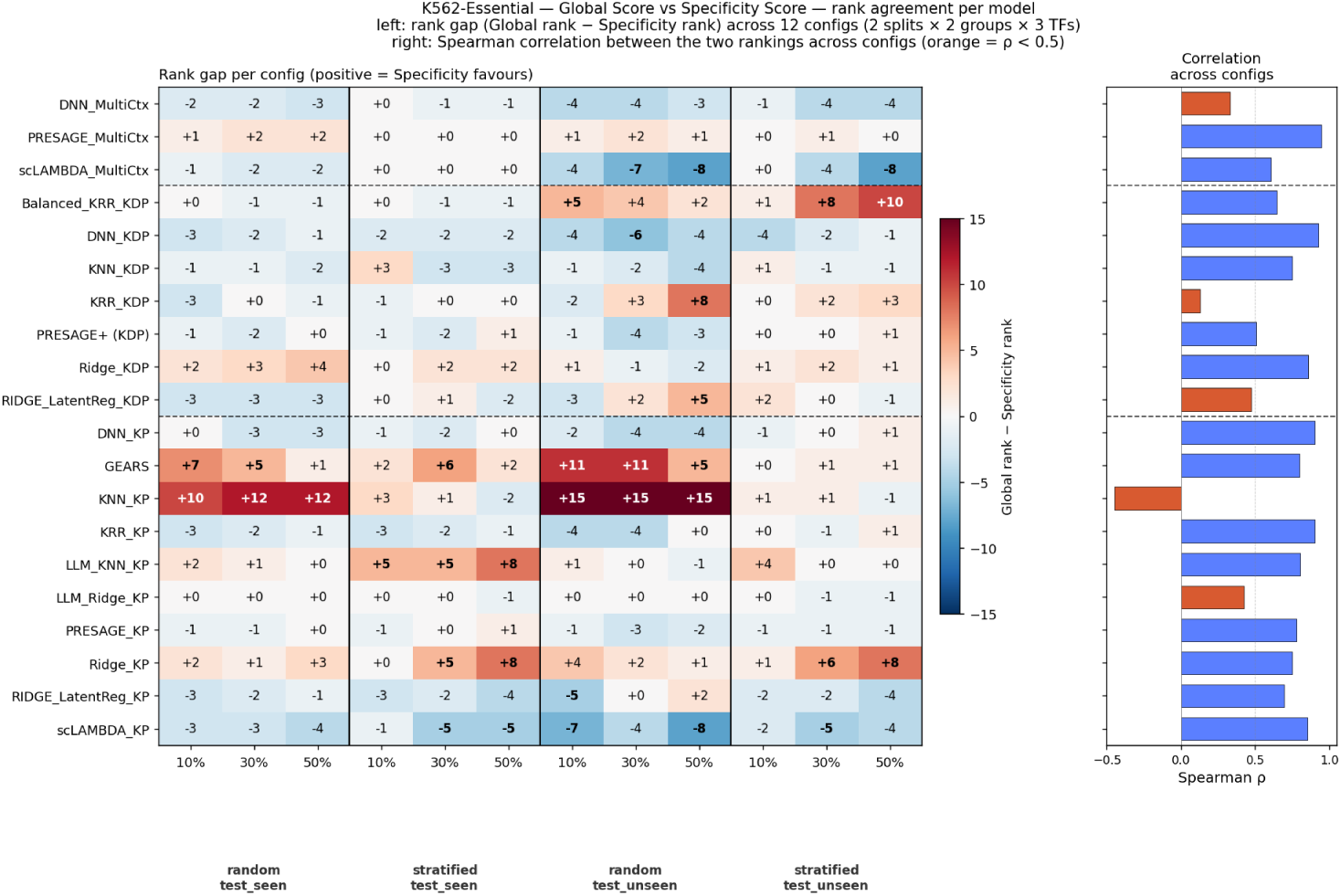
Agreement between Global and Specificity Score rankings per model (K562- Essential). Left: for each of the 12 configurations (random/stratified split seen/unseen perturbations training fractions of 10%, 30% and 50%), models were ranked by their Global Score and Specificity Score (averaged over random seeds; rank 1 = best), and each cell shows the rank gap (Global rank - Specificity rank). Each cell shows the rank gap, Global rank *−* Specificity rank (rank 1 = best). Positive values (red) mean the model is ranked better (closer to 1) by the Specificity Score than by the Global Score; negative values (blue) mean it is ranked better by the Global Score. Right: Spearman correlation between each model’s Global and Specificity ranks across the 12 configurations; orange bars mark <0.5. Dashed lines separate integration strategies (MultiCtx, KDP, KP).

This supports a core design principle of EL-PERTURB - ranking models by any single metric family can systematically favor particular architectural features. Therefore, we use the Global Score as our primary summary of model performance, with the Specificity Score and the per-metric breakdown as complementary views that reveal where a model’s strengths lie.

## 5 Discussion

The central question motivating EL-PERTURB is whether current genetic perturbation prediction models genuinely generalize beyond their training distribution, particularly under a data-limited setting - or whether their reported performance reflects increasingly sophisticated interpolation within a familiar neighborhood of the training data. Our results provide a clear and consistent answer: under rigorous stratified OOD evaluation, all tested models degrade substantially, architectural complexity confers no reliable advantage, and the fundamental bottleneck is not model design but the structure of the training data itself.

### The interpolation problem

Across all 20 model configurations, six architecture classes, and two dataset types, the pattern is the same: models perform relatively better under random train/test splits and fail under stratified OOD splits. This is the signature of interpolative models - systems that have learned to retrieve responses from a memorized neighborhood in perturbation representation space rather than to encode the underlying regulatory biology that generates those outcomes. The attention mechanisms of PRESAGE, the graph structure of GEARS, and the foundation model-derived features of scGPT and Geneformer all fail to provide meaningful protection against distributional shift. This is not a criticism of any individual architecture - it is a structural property of the entire class of perturbation prediction models currently in use. They are, in essence, sophisticated nearest-neighbor regressors. Moving the needle on OOD generalization will require approaches that go beyond perturbational similarity as the primary predictive signal - toward models that encode causal structure, the physical propagation of perturbation effects through regulatory networks, and the counterfactual logic of gene knockout-driven trans regulation.

### What the data tells us about model complexity

On essential gene screens - where cross-context response correlations are strong - multi-context MultiCtx models show a genuine advantage over single-context models at low training fractions, suggesting they can leverage shared response structure for domain adaptation. But this advantage disappears on genome-wide screens, where cross-context correlations are weaker, and simpler models including ridge regression, KNN, and LLM retrieval consistently outperform deep learning approaches. Taken together, these results argue against the field’s current trajectory of pursuing increasingly complex architectures as the primary path to better perturbation prediction. Model complexity is not the primary bottleneck.

### Foundation model embeddings

The finding that scGPT and Geneformer embeddings - including causally-inspired *in silico* perturbation embeddings - provide no improvement over curated knowledge priors deserves emphasis. This is consistent with earlier findings [9–11, 27] indicating that current transcriptomic foundation models do not yet encode perturbation-relevant biology that surpasses simpler alternatives. The path forward likely requires pretraining on large-scale perturbation atlases with learning objectives that explicitly model causal intervention rather than masked expression reconstruction from covariation patterns.

### Implications for future benchmarking

Our design tests varying levels of generalization - few-shot in-context learning, perturbation-exclusive (unseen) generalization, and multi-context learning. EL-PERTURB establishes several methodological standards we argue should be adopted as community norms. First, stratified OOD splitting based on functional clustering of response profiles should replace random splitting as the primary evaluation protocol. Second, few-shot training regimes (10–50% in-context coverage) should be the default evaluation setting. Third, multi-metric evaluation spanning error, correlation, rank-based discriminability, and manifold preservation is necessary to produce architecture-agnostic rankings. And fourth, model capability should be interpreted in light of the compositional diversity of the evaluation data used, reflected in the shared response structure both across perturbations in-context and across reference-target screens.

### Limitations

Our study focuses on few-shot single-gene perturbation effect prediction in a target cell line, and does not consider zero-shot generalization to altogether novel (unseen) cell lines. Our results do not imply a ceiling on perturbation prediction performance. Extensions that could improve on reported benchmarks include more stringent data quality control to handle technical and biological confounders, more extensive hyperparameter optimization, and incorporation of additional biological prior knowledge sources including multi-modal knowledge graphs; in particular, reliance on biological mechanism priors that mitigate ascertainment bias inherent in the biomedical literature will be needed to improve the representation of under-characterized genes. Loss functions better aligned with target metrics (such as Weighted MSE) can potentially improve the reported scores - although limited testing of this assumption leads to mixed results (we find that replacing MSE loss with WMSE improves the test WMSE, but this is countered by a *decrease* along the other metrics that similarly evaluate perturbation-specific discriminability). Our study is restricted to pseudobulk expression response prediction - models that directly leverage single-cell-resolution statistics may capture subtler cross-context correlations. We also note that conclusions are drawn from a specific set of publicly available Perturb-seq screens; results may differ on screens with different perturbation distributions, cell type compositions, or experimental platforms.

The deeper limitation - and the one that points most directly to what the field must build next - is that all evaluated models share a common architectural constraint: they predict by analogy, not by mechanism. Truly generalizable perturbation prediction will require models that encode the causal structure of regulatory biology, the context-specific protein-protein and transcriptional regulatory interactions that determine how perturbation effects propagate, and temporal data capturing the dynamic trajectory of cellular response rather than its static endpoint. Such mechanism-aware models, trained on large-scale perturbation atlases [3, 6] with causally-grounded learning objectives, represent the frontier that EL-PERTURB’s generalizability benchmark is designed to evaluate.

## 6 Conclusion

We introduced EL-PERTURB, a systematic benchmarking framework for out-of-distribution generalization in gene perturbation effect prediction. Across 20 model configurations spanning six architecture classes, evaluated along different axes of generalization on essential gene and genome-wide Perturb-seq screens, we find that all current perturbation prediction models fail under realistic distributional shift - and that architectural complexity, including the use of single-cell foundation model embeddings, provides no reliable remedy. These models are fundamentally interpolative: they retrieve responses by analogy to training examples rather than by encoding the causal biology that generates them.

EL-PERTURB establishes the first rigorously stratified OOD benchmark for perturbation prediction and contributes three methodological standards we propose the community adopt: stratified OOD splitting as the default evaluation protocol, few-shot training regimes as the default data setting, and multi-metric aggregation as the default scoring approach. Together, these standards provide the infrastructure needed to distinguish genuine progress in perturbation effect prediction from increasingly sophisticated interpolation - and to evaluate the causally-grounded, mechanism-aware models that the field must build next.

## 7 Data and Code Availability

This study makes use of publicly available datasets. All original data sources are referenced in the text. Python scripts needed to train and evaluate the ML models are shared at URL: https://github.com/ElucidataInc/EL-PERTURB.git. The complete evaluation results (individual metrics, global and specificity scores) across all benchmarking experiments conducted in the study are also shared in the GitHub repo.

## Supporting Information - Detailed Methods

### Datasets and Preprocessing

To evaluate model performance across diverse biological contexts, we integrated essential gene CRISPRi perturbation screens from four distinct human cell lines: K562, HepG2, RPE1, and Jurkat [22, 23]. Raw scRNA-seq profiles were normalized to a fixed target sum and log-transformed. To retain high-confidence perturbation measurements, each screen was filtered in three sequential steps using the cell-load package [28]. First, individual cells were retained only if they showed at least 50% knockdown of their target gene relative to non-targeting controls. Second, target genes were retained only if the mean knockdown efficiency across filtered cells exceeded 50%. Third, target genes with fewer than 20 cells remaining after the preceding filters were excluded to ensure sufficient coverage for pseudobulking. Highly variable genes (HVGs) were identified independently for each cell line, and the final feature set was defined as the union of common HVGs and common target genes perturbed across all four cell lines. Following feature selection, data underwent a second round of normalization and log-transformation to provide a stabilized scale for model training. Ultimately, each dataset comprised 1060 shared target gene perturbations measured over a common feature space of 1463 highly variable genes. The primary prediction target was the pseudobulk transcriptomic response induced by single-gene CRISPR knockdowns. For each gene knockdown, the transcriptional response was computed as the shift from the mean log-normalized expression of non-targeting control cells, yielding a delta-expression vector that served as the ground truth for both training and evaluation.

To assess generalizability beyond essential gene screens, we applied the same preprocessing and splitting pipeline to single-cell data from four genome-wide perturbation screens in K562, HCT116 and HEK293 cell lines, and resting CD4+ T cells [22, 24, 25]. The only modification was at the quality control step, where per-cell and per-gene knockdown thresholds were relaxed from 50% to 30% to accommodate the lower knockdown efficiencies in the genome-wide screens. Ultimately, each genome-wide screen comprised 3065 shared target gene perturbations measured over a common feature space of 3202 highly variable genes.

### Modeling Framework

Perturbation effect prediction was framed as a supervised regression task: given an input representation vector I*_P_* encoding information about gene perturbation P, a model f learns to predict the delta-expression response Δ = f(I*_P_* ) in a target cell line. Input representations combined two complementary sources of information: biological prior knowledge about the function and molecular context of the target gene, and experimental Perturb-seq response profiles of the same gene measured in other available cell lines (reference contexts), which capture transferable response signatures.

We evaluated three integration strategies forming the core axes of our experimental design:

**Knowledge Priors only (KP):** Single-context models trained exclusively on in-context data using knowledge-based perturbation embeddings as input features, with no access to cross-context experimental data.

**Knowledge + Data Priors (KDP):** Single-context models trained on in-context data, augmented with cross-context Perturb-seq response profiles as additional input features alongside knowledge embeddings. Cross-context data informs the input representation but is not used as training labels.

**Knowledge + Data Priors + Multi-context Training (MultiCtx):** Multi-context models trained jointly on target and reference cell line data in a context-aware manner, with cell line identity provided as an additional input. This configuration allows the model to learn shared perturbation response structures across contexts while being evaluated on its ability to predict target-context responses.

All models received the same fixed knowledge-based perturbation embeddings adopted from the PRESAGE publication: a concatenation of embeddings from 40 heterogeneous knowledge sources spanning curated functional databases (GO, Reactome, STRING), large-scale CRISPR screening data (DepMap), and pretrained large language models (ESM, BioGPT, GenePT)[13]. Each source-specific embedding was reduced to a fixed dimension of d=128 via PCA prior to concatenation, with zero-padding applied where source dimensionality was lower than 128. All models thus receive the same effective input, ensuring fair comparison across architectures.

We benchmarked six architecture classes spanning a range of model complexity:

*Attention-based models:* PRESAGE [13], an attention-based deep learning model integrating multi-source gene embeddings through dual pooling - a global softmax-weighted average combined with gene-specific graph attention convolution (GATConv). Evaluated in three configurations: standard single-context (KP), cross-context feature augmentation (KDP, referred to as PRESAGE+), and multi-context training (MultiCtx). For the MultiCtx variant, context identity was injected via 128- dimensional cell-type embeddings derived from control cell populations, with feature selection based on ANOVA F-scores across contexts. See the next subsection for additional details on PRESAGE implementation.

*Multi-layer neural networks:* A feedforward MLP with four hidden layers and Feature-wise Linear Modulation (FiLM) [29] for context injection, evaluated in both single-context (KP, KDP) and multi-context (MultiCtx) settings. Layer normalization and dropout were applied for regularization.

*Linear and kernel methods:* L2-regularized ridge regression (KP, KDP), kernel ridge regression (KRR) with linear and RBF kernels using a multi-source combined kernel matrix (KP, KDP), a balanced KRR variant with equal weighting of knowledge and data kernels (KDP only), and a latent-ridge model using SVD of the combined kernel matrix as input features to ridge regression (KP, KDP), and K-nearest neighbors (KNN) regression using the combined kernel matrix as a distance measure (KP, KDP).

*LLM-based models:*LangPert [26], which uses LLM evidence synthesis (OpenAI GPT-5.1) to assign nearest training-set neighbors to each test perturbation followed by KNN aggregation (KP only); and a Ridge-LLM variant using LLM-generated pathway activation scores as per-perturbation input features to L2-regularized linear regression (KP only). Refer to SI Table S4 for the LLM prompts used. In internal evaluation, we found context-agnostic prompts without specific reference to the target cellular context yielded relatively better predictions.

*Additional deep learning baselines:* We have included two reference models: GEARS [12], a graph neural network approach combining a gene-gene co-expression graph with a prior knowledge graph (KP only); and scLAMBDA [19], a variational autoencoder framework that leverages LLM-derived gene embeddings to disentangle basal cell states from perturbation-specific transcriptional responses (KP, MultiCtx). To enable a fair comparison, both GEARS and scLAMBDA were trained using the PRESAGE perturbation embeddings (40 sources concatenated) as input featurization. In GEARS, the original Gene Ontology-based graph was replaced by a gene-gene similarity graph grounded in the PRESAGE gene embeddings.

Our intent was to include representative model classes of varying complexity, with model selection driven by alignment with the evaluation framework we propose. A core thesis of our work is that existing studies assess models in restrictive settings. Our design tests varying levels of generalization, and we have included published models evaluable under these settings. CPA, scGen and STATE are applicable only under context transfer (i.e., for seen perturbations profiled in other cell lines) and do not generalize to the unseen-perturbation setting, and were therefore excluded. TxPert could not be included on reproducibility grounds - its public release exposes only inference with pretrained checkpoints (covering only a subset of the 40 knowledge sources/graphs used by PRESAGE), with no training entry point, and the best-performing configurations rely on proprietary knowledge graphs that have not been released.

The complete list of model configurations and hyperparameter ranges is provided in SI Table S1. All models except PRESAGE variants were tuned via five-fold cross-validation on the in-context training data, using MSE as both the training and validation loss. For PRESAGE, we retained the optimized hyperparameter configurations reported by the original authors [13]. Five-fold cross-validation was used solely to determine the optimal number of training epochs, as the original train/test splits do not include a dedicated validation set. Specifically, we recorded the epoch at which the lowest validation loss was achieved across each fold and used the maximum best epoch count to set the training duration for final model fitting.

### Additional Details on PRESAGE Model Variants

PRESAGE (Perturbation Response EStimation with Aggregated Gene Embeddings), is an attention-based deep learning framework designed to predict transcriptomic changes from high-dimensional target gene embeddings [13]. The architecture utilizes target gene representations derived from 40 heterogeneous knowledge sources, spanning curated functional databases such as GO, Reactome, and STRING; large-scale CRISPR screening data from DepMap; and embeddings from pretrained biological foundation models including ESM, BioGPT, and GenePT. These source-specific embeddings are integrated through a dual-pooling mechanism that combines a global pooling step—learning dataset-wide source weights via a softmax-weighted average—with a gene-specific attention pooling step utilizing graph attention convolution (GATConv). An attention hyperparameter controls the final weighted average of these pooled vectors, which is then mapped to predicted log-fold changes through a linear transformation.

To evaluate the impact of knowledge and experimental data priors, we implemented three distinct configurations of the PRESAGE architecture:

**Standard PRESAGE**: This configuration was trained strictly on target context perturbation data using the 40 biological knowledge sources as input features.

**PRESAGE+ (PPLUS)**: This variant incorporated cross-context Perturb-seq data from reference contexts as indirect input features. Per-gene response profiles from reference screens were compressed into latent embeddings via Principal Component Analysis (PCA) and added to the model as an additional knowledge source. While the training outcomes remained restricted to the target context, the model utilized these reference-derived response profiles to featurize the input perturbations.

**Presage_MultiCtx**: This multi-context variant was trained directly on merged perturbation response data from both target and reference contexts. To differentiate between identical perturbations across biological environments, perturbations in the merged dataset were encoded as a combination of the target gene and cell-type identity (ex. k562:BMAL1, rpe1:BMAL1). Thus, for each target gene, the final embeddings input to the model included the 40 source-specific knowledge embeddings plus an additional cell type identity embedding to explicitly account for context-specific baseline shifts while learning shared perturbation response patterns through direct supervision. The cell type identity embeddings were 128-dimensional cell-type embeddings (matching the knowledge source embedding size) derived from the control cell populations of each context. Feature selection for these embeddings was performed by identifying the top 128 genes with the highest ANOVA F-scores across contexts, ensuring the capture of discriminative baseline transcriptomic states.

### Train/Test Splitting and the Few-Shot OOD Evaluation Framework

A central methodological contribution of EL-PERTURB is its train/test splitting strategy, designed to emulate the realistic few-shot prediction setting and enable rigorous quantification of OOD generalization. We employed a leave-one-context-out design in which one cell line served as the target context, with the remaining cell lines providing reference data.

To emulate realistic data scarcity, in-context training fractions (TF) were set at 10%, 30% and 50% of the target screen perturbations - corresponding to approximately 100–500 perturbations for essential gene panels and 300–1,500 perturbations for genome-wide screens. Held-out test perturbations were partitioned into two subsets. The *seen* subset comprises perturbations withheld from the target training set but profiled in reference contexts, specifically probing cross-context transfer. The *unseen* subset - fixed at 10% of total perturbations and strictly removed from all reference contexts - comprises genes absent from all training data, probing genuine domain adaptation and extrapolation to novel biological programs.

To evaluate the impact of distributional shift on model performance, we contrasted two splitting strategies. The *random split* draws train and test perturbations from a shared distribution in transcriptional response space, testing interpolative performance. The *stratified OOD split* applies K-means clustering to ground truth pseudobulk response profiles and assigns entire functional clusters to either train or test, ensuring that train and test sets occupy functionally distinct regions of transcriptional space. The OOD generalization gap for each model is then quantified as the percentage change in performance between the random and stratified conditions, providing a direct and comparable measure of distributional robustness across all the model configurations.

### Cluster-Based Strategy for Data Partitioning

To model the OOD setting, we group and segregate perturbations by their ground-truth response profiles. Perturbational expression response is a direct proxy for the functional role of a gene, since its inactivation largely impacts the biological programs it participates in or regulates. Cluster-wise sorting of response vectors therefore simulates functional distribution shift between folds, ensuring genes with similar cellular roles are assigned together to training or held out together for testing, avoiding leakage. We chose this over splitting by gene function or family, since gene-to-phenotype relations are complex, context-dependent, and not accurately reflected by molecular-level functions. Protein kinases, for example, share structural/biochemical attributes but participate in diverse pathways and may have widely varying knockdown impacts; conversely, proteins with distinct molecular activities can produce similar post-knockdown phenotypes if they act within a shared program. Segregation by biochemical properties is therefore expected to be unreliable for OOD splits given pleiotropy and the disparity between molecular- and cellular-scale impact; this is better captured by genome-wide knockdown response.

Nonetheless, we have evaluated the resulting distribution shift in the prior-knowledge (PRESAGE) gene embedding space as well. Comparing train/test separation under random versus stratified splitting in the input gene embedding PCA space (SI, Table S5), the stratified splits generate more separated folds even in this independent representational space, validating our strategy.

To characterize the biological composition of the partitions more directly, we also ran pathway enrichment (Enrichr [30]; GO Biological Process 2023, Reactome 2022, MSigDB Hallmark 2020) on the train, test-seen and test-unseen gene sets for every configuration (train fractions 10/30/50%, three seeds, both split types), using the screened target genes as background so that enrichment reflects the split rather than the biased selection of screened genes. Under stratified splitting, all partitions were strongly pathway-enriched (SI, Table S6); under a size-matched random split, no partition exceeded 0.3 significant terms at any training fraction. The stratified partitions are also enriched for near-disjoint programs: mean pairwise Jaccard overlap between partitions’ enriched-term sets was 0.008, 0.000 and 0.001 at TF10/30/50, respectively. The stratified split therefore produces partitions that are each biologically coherent and mutually distinct at the pathway level, whereas random splitting yields essentially no enrichment structure.

Choice of K: SI, Figure S6 shows the dependence of the separation between the train and test splits as a function of the number of transcriptomic clusters. The degree of distribution shift is tuned by the number of clusters (K), with the splits segregating more clearly at larger cluster sizes and approximating random sorting in the large K limit. We generated results for stratified splits obtained with K=10, so our random and stratified partitions represent two ends of a spectrum.

### Evaluation Framework

Model predictions were evaluated against ground truth pseudobulk delta-expression profiles on held-out perturbations in the target context using eight complementary metrics organized across three levels of biological specificity.

*Transcriptome-level accuracy* was assessed using **Mean Squared Error (MSE)** and **Pearson correlation** across all genes, providing global sanity checks on prediction magnitude and direction.

*Biologically significant response accuracy* was measured using Pearson correlation restricted to the top 50 differentially expressed genes (DEGs) of each perturbation (**Pearson-TopK**), focusing evaluation on the genes most strongly affected by each perturbation.

*Perturbation-specific discriminability* - the most stringent level - was assessed using four metrics designed to penalize mode collapse and reward the capture of perturbation-unique signals: **Weighted Mean Squared Error** (WMSE, with gene weights computed as in Mejia et al. [8]); **Pearson- Pert** (correlation of predicted shift relative to the mean of all training perturbations rather than the non-targeting control, isolating perturbation-specific signal); two retrieval-based **Rank Scores (L1 and Cosine)**, equivalent to the transposed rank metric in PerturBench [5] and the Perturbation Discrimination Score used in the ARC Virtual Cell Challenge [31]; and a **Matrix Distance** metric - the Frobenius norm of the difference between predicted and true pairwise cosine similarity matrices - to assess global manifold structure preservation. The perturbed gene was excluded from all metric calculations to focus evaluation on trans-effects, ignoring the trivial knockdown-efficacy signal.

All raw metrics were converted to Scaled Scores normalized against a Cell-Mean (CM) baseline - the mean response profile across all training perturbations - representing the mode-collapse reference:

**For correlations and rank scores** – Scaled Score = (Model Score *−* Baseline Score) / (1.0 *−* Baseline Score)

**For error and distance metrics** – Scaled Score = (Baseline Score *−* Model Score) / Baseline Score

A Scaled Score of 1.0 represents perfect prediction; 0.0 represents performance equal to the CM baseline; negative values indicate performance worse than the naive baseline.

Two meta-scores were derived from the eight scaled metrics. The *Global Score (GS)* is the arithmetic mean of all eight scaled metrics, providing a rounded assessment across magnitude, correlation, retrieval, and manifold preservation, computed separately on seen and unseen test subsets. The *Specificity Score (SS)* is the mean of the five perturbation-specific metrics (WMSE, Pearson-Pert, Matrix Distance, Rank-L1, Rank-Cosine), specifically evaluating a model’s capacity for genuine discrimination among distinct perturbations rather than memorization of shared transcriptional shifts.

### Compute Resources

Given the combinatorial experimental design - 3 training fractions *×* 2 splitting strategies *×* 3 random seeds - a total of 18 configurations were evaluated per model variant. Model training, hyperparameter optimization and evaluation of PRESAGE, GEARS, scLAMBDA and DNN was performed on NVIDIA L4 GPUs. All other models were trained and evaluated on a MacBook CPU machine (Apple M1, 16 GB RAM).

## Supporting Information - Tables

**Table S1:** Full set of model configurations (model type. *×* **input featurization) evaluated**, along with the specific hyperparameters associated with each model type which were tuned via five-fold CV on the in-context training data.

| Model type | Configurations (input features + training contexts) | Model architecture and Training hyperparameters (ranges) |
| --- | --- | --- |
| **PRESAGE** | KP, KDP (Presage+), MultiCtx | Refer to SI Detailed Methods (“*Additional Background on PRESAGE Model Variants*”) for details |
| **DNN (MLP)** | KP, KDP (single-context, concatenates the knowledge embeddings and the cross-context response profiles), MultiCtx | MLP feedforward neural network (4 hidden layers) with FiLM-type context modulation at hidden layer #2; LayerNorm and dropout based regularization; fixed learning rate = $1e-3$; max_epochs = 100<br>- Input dimension: 128, 512<br>- Hidden layer dimension: 128, 512<br>- Dropout rate: 0, 0.2, 0.5<br>- Batch size: 16, 32, 64<br>- Target context data fraction per training batch: *None* (random sampling), 0.25 |
| **Ridge Regression** | KP, KDP | L2-regularized linear regression<br>- Alpha (regularization parameter): 0.01 - 1000 |
| **Kernel Ridge Regression (KRR)** | KP, KDP | Multi-source combined kernel matrix (each individual input source weighted equally)<br>- Kernel type: Linear, RBF<br>- Alpha (regularization parameter): 0.01 - 10<br>- Gamma (bandwidth, applicable to RBF kernel only): 0.001 - 0.1 |
| **Balanced_KRR** | KDP only | Aggregated kernels obtained from knowledge embeddings and cross-context response data are weighted equally<br>- Kernel type: Linear, RBF<br>- Alpha (regularization parameter): 0.01 - 10<br>- Gamma (bandwidth, applicable to RBF kernel only): 0.001 - 0.1 |
| **K-Nearest Neighbors Regression** | KP, KDP | Mean of k-nearest neighbors in the embedding space (the combined kernel specifies the distance matrix)<br>- # of nearest neighbors (k): 1 - 50 |
| **Ridge_LatentReg** | KP, KDP | Latent perturbation embeddings obtained from SVD on the combined kernel matrix, used as input features for L2-regularized regression<br>- Alpha (regularization parameter): 0.1 - 1000 |
| **LLM_KNN**<br>(adpated from LangPert, Martens et al., 2025) | KP only (single-context training) | LLM retrieval of k nearest training set neighbors (up to max 5) for each test perturbation, followed by k-NN aggregation<br>- No tunable hyperparameters<br>- OpenAI model: `gpt-5.1-2025-11-13`; effort='medium' |
| **LLM_Ridge** | KP only (single-context training) | LLM prompting to obtain the predicted activation of each of a set of cellular pathways (186 KEGG gene sets from MSigDB) given a gene perturbation, which defines the input feature vector used in L2-regularized regression<br>- Alpha (regularization parameter): 0.01 - 1000<br>- OpenAI model: `gpt-5.1-2025-11-13`; effort='medium' |
| **GEARS**<br>(Roohani et al., 2023) | KP only (single-context training) | Graph-based geometric deep learning, combines a context-specific gene-gene co-expression graph (constructed from the non-targeting control cells) and a knowledge graph based on gene annotation similarity between perturbations<br>- max_epochs: 20 |
| **scLAMBDA**<br>(Wang et al., 2024) | KP, MultiCtx | Deep generative variational autoencoder that predicts context-specific responses conditioned on prior knowledge gene embeddings and the unperturbed cell state<br>- training_epochs: 100<br>- training_batch_size: 500<br>- step_size: 30<br>- gamma: 0.2<br>- learning_rate: $5e-4$<br>- Validation fraction: 20% of training data (used to select and save the best checkpoint) |

**Table S2:**
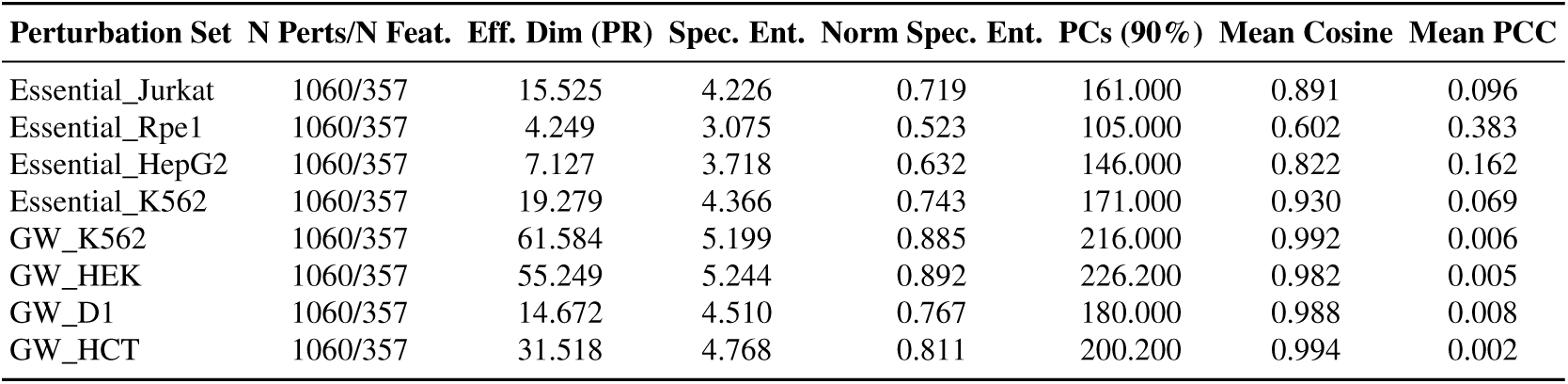
Comparison of perturbation diversity and effective dimensionality of response programs across different cell lines and perturbation sets. For genome-wide screens, numbers represent the mean over five randomly downsampled submatrices. N Perts: No. of sampled perturbations; N Feat.: No. of gene expression features; Eff. Dim.: Effective Dimensionality/Participation Ratio from SVD eigenspectrum; Spec. Ent.: Spectral Entropy; Norm. Spec. Ent.: Normalized Spectral Entropy; PCs (90%): No. of PCs explaining 90% of the response variance; Mean Cosine: Mean pairwise Cosine Distance between perturbation responses; Mean PCC: Mean pairwise Pearson Correlation between perturbation responses.

**Table S3:**
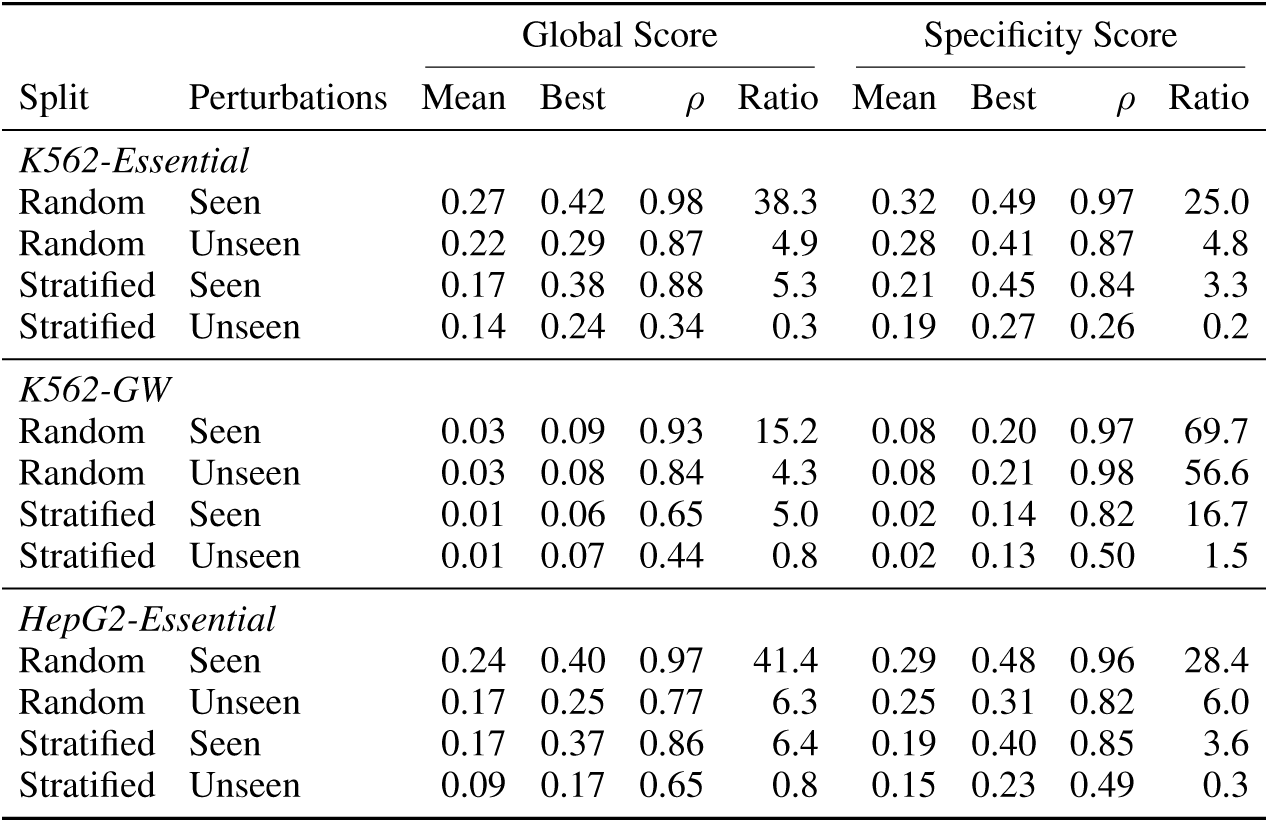
Reproducibility of model rankings across random seeds. For each dataset, score and evaluation condition (random/stratified split *×* seen/unseen perturbations), we report the mean score across all models and runs, the best model’s mean score (averaged over training fractions and seeds), the between-seed rank correlation, and the between-model to within-model variance ratio. The between-seed rank correlation is the Spearman correlation between model rankings obtained with different random seeds at the same training fraction, averaged over all seed pairs and training fractions. The variance ratio compares the variance of the models’ mean scores with the average variance of each model’s score across seeds; values below 1 indicate that models differ from each other less than a single model varies across seeds. Mean: mean score across models and runs; Best: best model’s mean score; *ρ*: between-seed Spearman correlation; Ratio: between-model to within-model variance ratio.

**Table S4:** Prompts used for LLM-based retrieval of perturbation predictions.

| Model | Input Prompt |
| --- | --- |
| **LLM-based KNN Aggregation** | Task: You are given a list of human query genes. For EACH QUERY GENE, choose ONE TO FIVE genes from the CANDIDATE LIST that are MOST SIMILAR in terms of perturbation response.<br>Similarity definition: Knockout/knockdown causes similar transcriptome-wide expression changes.<br>Base your reasoning on evidence from:<br>- Shared pathways, complexes, protein or TF interactions<br>- Downstream processes impacted<br>- Functional genomics screens and phenotype similarity<br>DO NOT restrict context to any particular cell-line or type. Use all available human biological information.<br>Rules:<br>- Each query gene MUST MAP TO AT MOST 5 candidate genes from the provided list<br>- Use ONLY the provided candidate list for selection<br>- Do not return any reasoning, only return the list of selected genes for each query<br>Query genes: [genes]<br>Candidate gene list: [candidates] |
| **LLM-based Ridge Regression** | Task: You are a functional genomics expert predicting pathway-level effects of human gene [gene] KO/KD.<br>Return EXACTLY n floats in [-1, 1], one per pathway below, in the same order.<br>Rules:<br>- -1 = strong inhibition of that pathway<br>- +1 = strong activation of that pathway<br>- 0 = no expected effect<br>- Use biological reasoning; be deterministic.<br>- No explanations. Strict structured output only.<br>- Never omit entries. If unsure, use 0.0.<br>Query Gene: [gene]<br>Pathway chunk: [n pathways with name and 5 sample genes per pathway] |

**Table S5:** Train/test-seen/test-unseen separation in PCA space of PRESAGE gene embeddings (40 sources concatenated), averaged over seeds, for random/stratified splits (target = K562-Essential).

| TF | kNN purity (rand/strat) | Calinski-Harabasz (rand/strat) | Silhouette (rand/strat) |
| --- | --- | --- | --- |
| 10 | 0.637 / 0.719 | 1.21 / 9.95 | 0.001 / 0.023 |
| 30 | 0.433 / 0.520 | 1.32 / 8.21 | -0.004 / -0.005 |
| 50 | 0.417 / 0.500 | 1.27 / 6.63 | -0.004 / -0.004 |

**Table S6:** Mean number of significantly enriched functional annotation terms (adjusted p < 0.05) per partition under random/stratified splitting. , averaged over seeds (target = K562-Essential).

| Partition | TF10 (rand/strat) | TF30 (rand/strat) | TF50 (rand/strat) |
| --- | --- | --- | --- |
| Train | 0 / 47.7 | 0.3 / 21.0 | 0 / 16.0 |
| Test Seen | 0 / 13.0 | 0 / 8.7 | 0 / 40.7 |
| Test Unseen | 0.3 / 89.3 | 0.3 / 89.3 | 0.3 / 89.3 |

## Supporting Information - Figures

**Figure S1:**
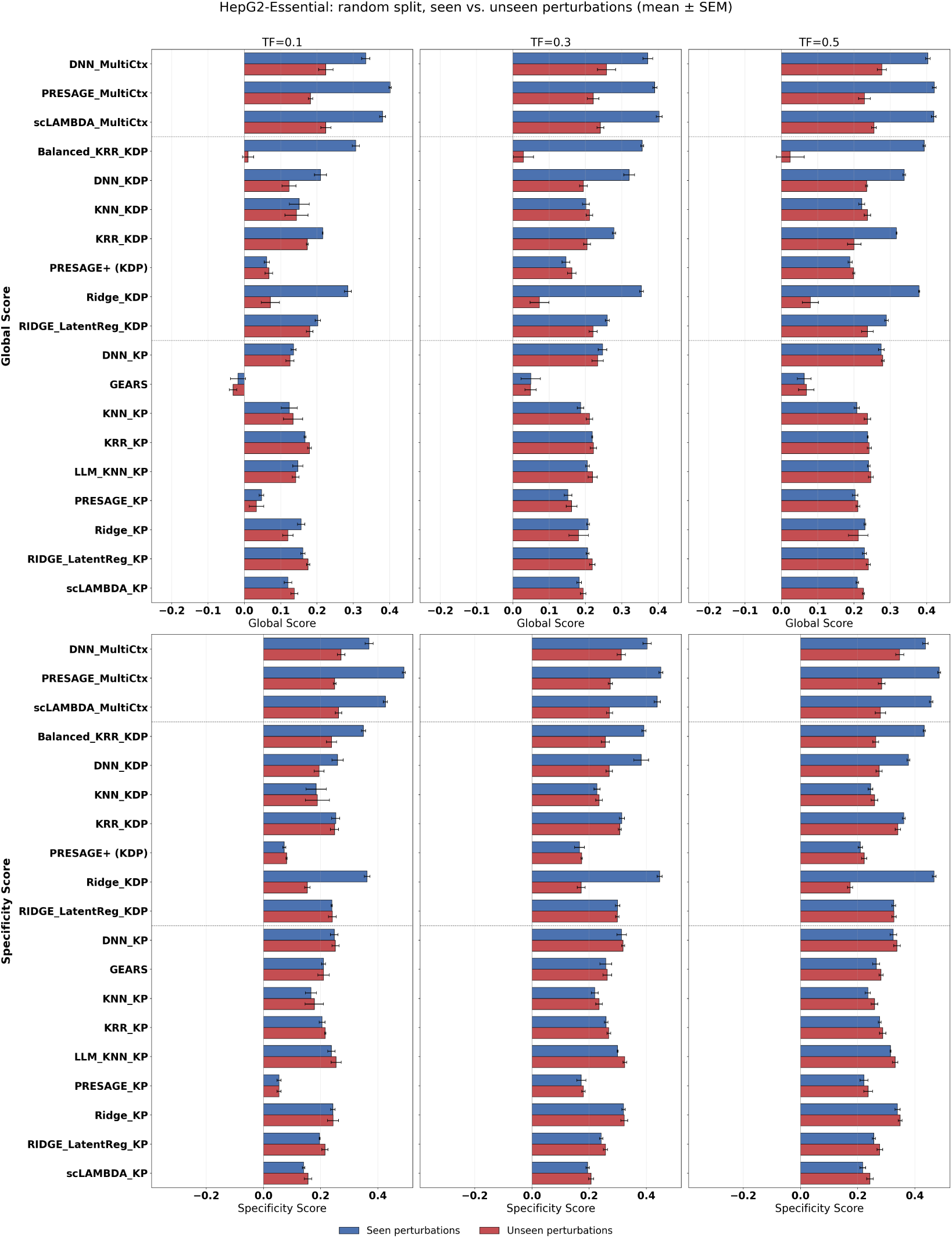
Model performance under random data splitting on seen vs. unseen perturbations (essential gene screen; target context = HepG2). Grouped bar plots compare performance on seen (blue) and unseen (red) perturbations under the random split. Rows show the Global Score (top) and Specificity Score (bottom); columns show training fractions of 10%, 30% and 50%. Bars show the mean *±* SEM across random seeds.

**Figure S2:**
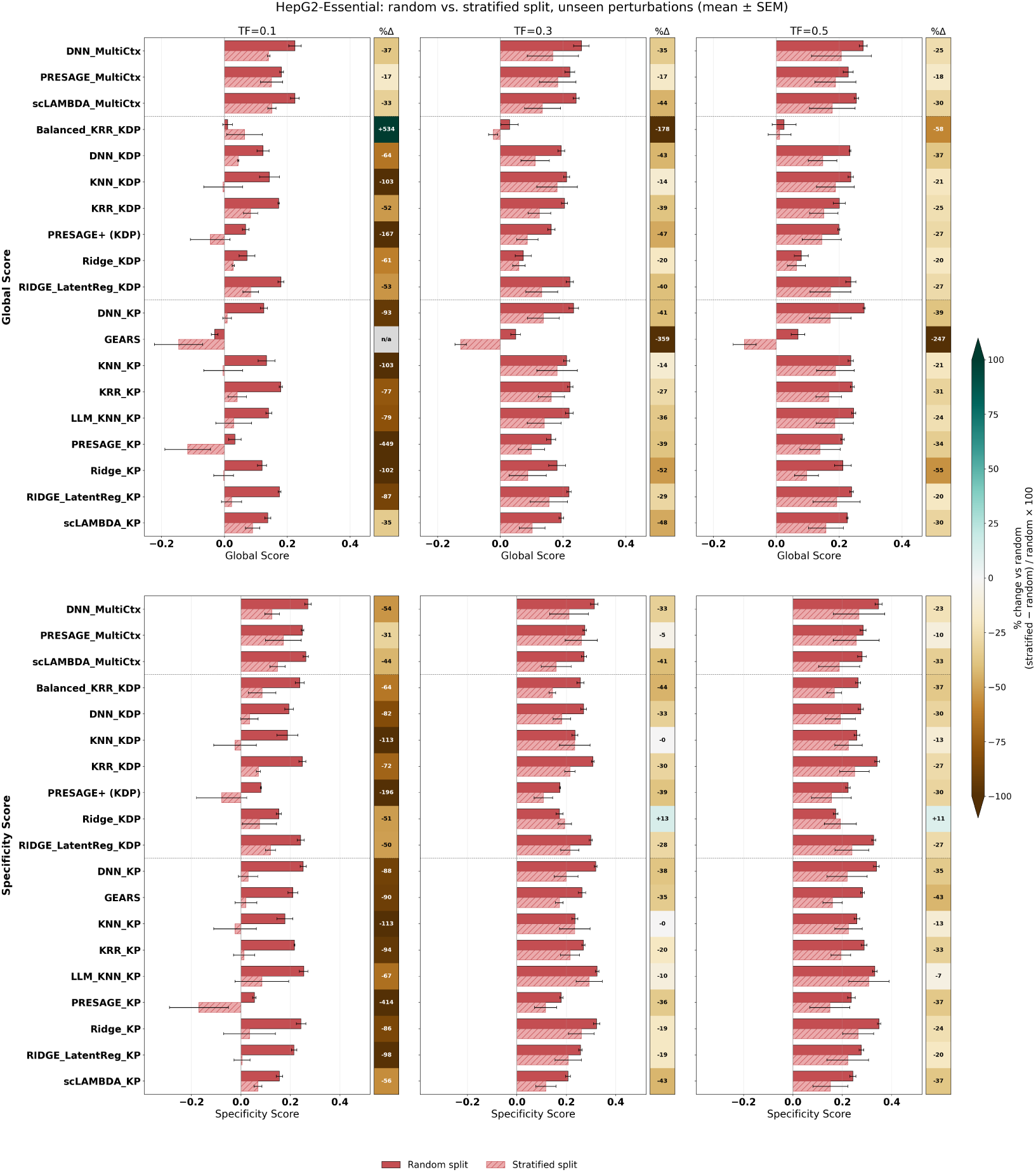
OOD generalization gap across models for essential gene screens (target context = HepG2). Grouped bar plots comparing performance on unseen perturbations under random (solid red) and stratified (light red, hatched) splits. Rows show the Global Score (top) and Specificity Score (bottom); columns show training fractions of 10%, 30% and 50%. Bars show the mean *±* SEM across random seeds. The column of squares to the right of each panel (%Δ) gives the relative change under the stratified split, (stratified *−* random) / random *×* 100, for each model; brown indicates lower and teal higher performance than under the random split, with the colour scale capped at *±*100%. Any cells marked n/a are omitted because the random-split score is below 0.01, where a relative change is not meaningful.

**Figure S3:**
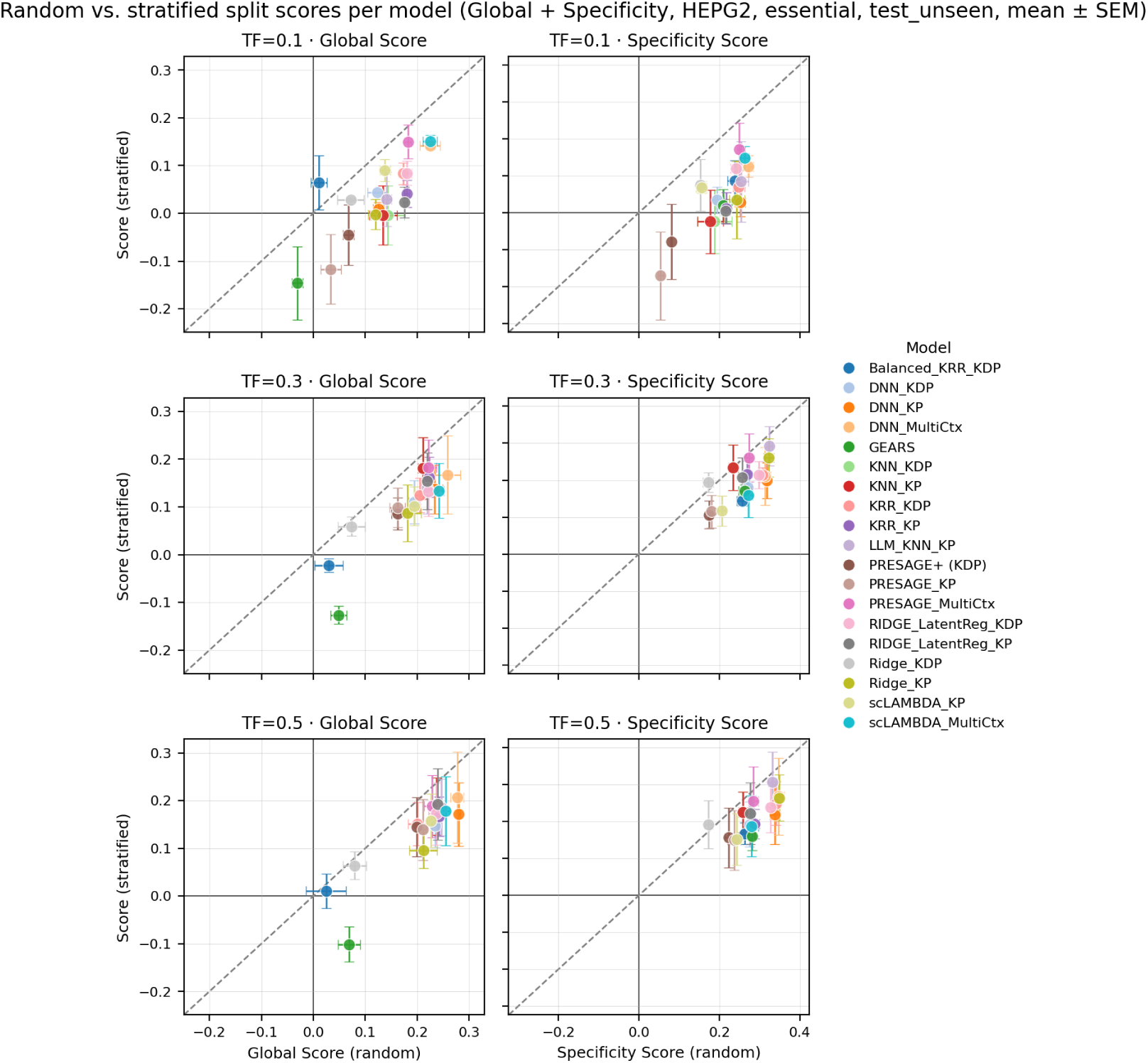
Aggregated view of regression performance for random vs. stratified splits across all model configurations on the HepG2-Essential panel. The OOD robustness of each model is reflected in its location on the 2D scatter plot relative to the diagonal. Each point represents the mean + error bar of prediction performance (Global/Specificity Scores) on the test-unseen set for one model configuration. TF: Training Fraction.

**Figure S4:**
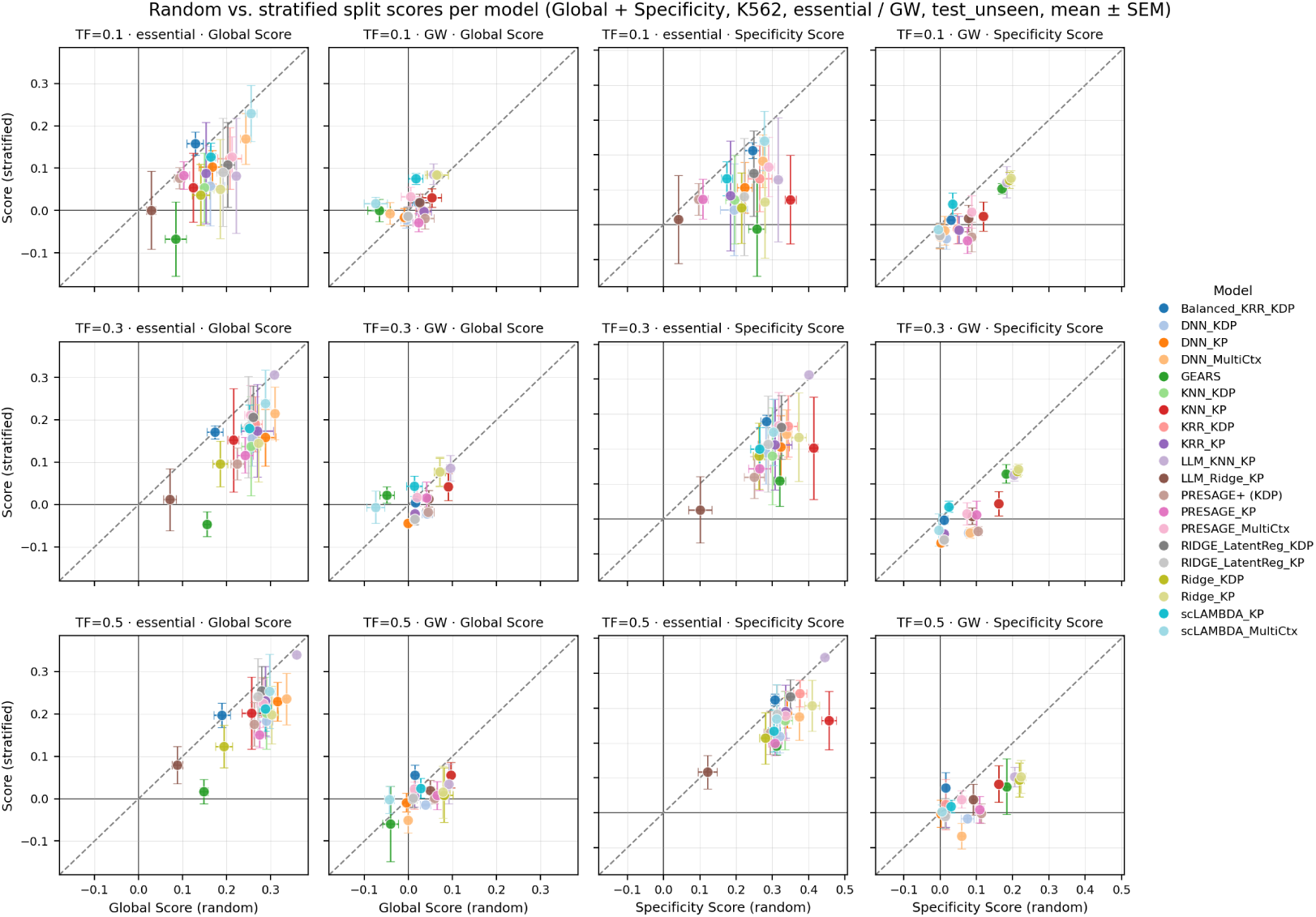
Aggregated view of regression performance for random vs. stratified splits across all model configurations on the K562 cell line. The OOD robustness of each model is reflected in its location on the 2D scatter plot relative to the diagonal. Each point represents the mean + error bar of prediction performance (Global/Specificity Scores) on the test-unseen set for one model configuration. Results for the essential-gene and genome-wide perturbation sets are displayed side by side, highlighting the general performance drop on the genome-wide panel relative to essential genes. TF: Training Fraction.

**Figure S5:**
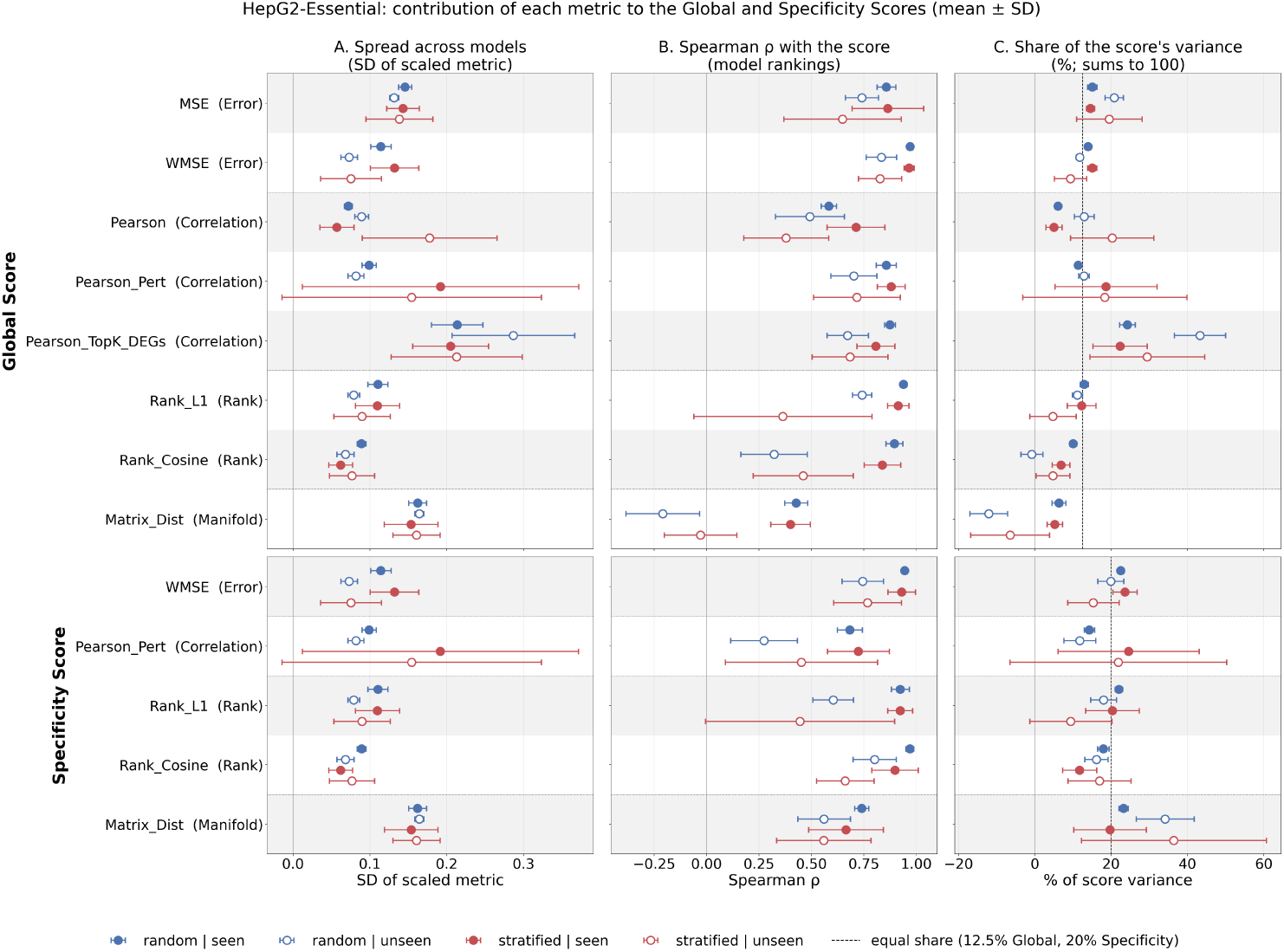
Effective contribution of each metric to the Global and Specificity Scores (HepG2- Essential). Rows show the Global Score (top; 8 metrics) and Specificity Score (bottom; 5 metrics). For each evaluation condition (blue = random, red = stratified split; filled = seen, open = unseen perturbations), we show (A) the spread of each scaled metric across models (standard deviation), (B) the Spearman rank correlation between each metric and the score across models (+1 = the metric ranks models in the same order as the score), and (C) each metric’s share of the score’s variance across models. Shares sum to 100% and can be negative when a metric works against the score; the dashed line marks an equal share (12.5% for the Global Score, 20% for the Specificity Score). Points show the mean *±* SD over the nine runs per condition (three training fractions three random seeds). Dotted lines separate metric families.

**Figure S6:**
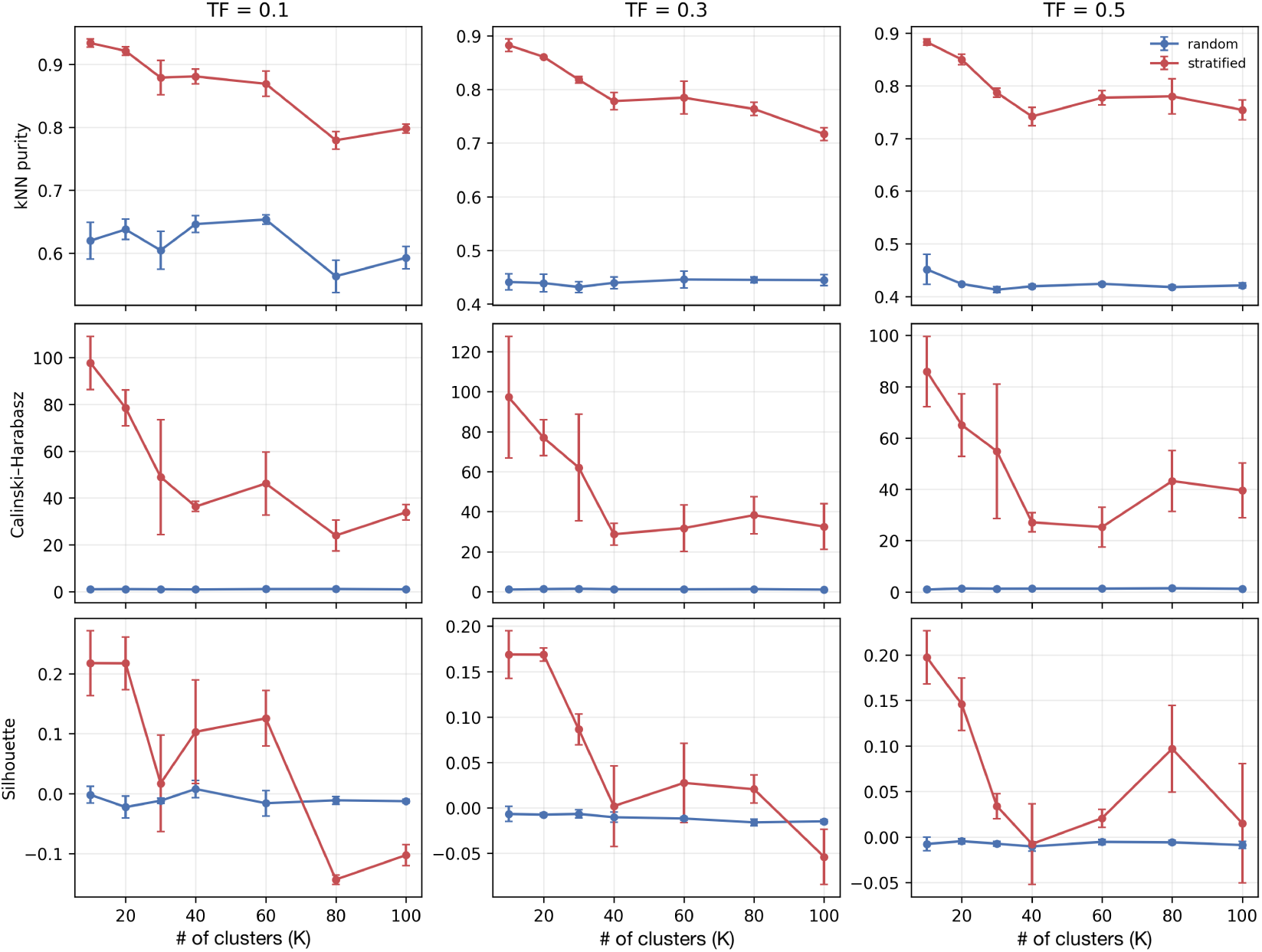
Segregation of K562-essential screen transcriptome-wide responses across train, test- seen and test-unseen splits as a function of the number of K-means partitions. The separation is quantified in terms of three metrics computed on the PCA-reduced ground-truth expression readouts (n_pca=50); error bars denote scatter across five random seeds. TF: Training Fraction.

## Notes

https://github.com/ElucidataInc/EL-PERTURB.git

### Competing Interest Statement

The authors have declared no competing interest.

